# scBFA: modeling detection patterns to mitigate technical noise in large-scale single cell genomics data

**DOI:** 10.1101/454629

**Authors:** Ruoxin Li, Gerald Quon

## Abstract

Technical variation in feature measurements such as gene expression and locus accessibility is a key challenge of large-scale single cell genomic datasets. We show that this technical variation in both scRNA-seq and scATAC-seq datasets can be mitigated by performing analysis on feature detection patterns alone and ignoring feature quantification measurements. This result holds when datasets have low detection noise relative to quantification noise. We demonstrate state-of-the-art performance of detection pattern models using our new framework, scBFA, for both cell type identification and trajectory inference. Performance gains can also be realized in one line of R code in existing pipelines.

## Background

Single cell genomics technologies have become a widely used technique for interrogating diverse problems related to gene regulation, including the identification of novel cell types and their regulatory signatures, trajectory inference for the analysis of continuous processes such as differentiation, high resolution analysis of transcriptional dynamics and characterization of transcriptional heterogeneity within populations of cells^1^. Of the different modalities that can be profiled, single cell RNA sequencing (scRNA-seq) is currently the most mature, and diverse scRNA-seq technologies are now available to cater towards specific applications. For instance, droplet-based methods such as Drop-seq currently have some of the highest throughput capture of cells and is suitable for rare cell type identification and characterization of tissue heterogeneity. On the other hand, so-called full-length transcript methods are able to measure alternative splicing and sequence individual cells more deeply, with the limitation of typically sequencing fewer cells.

scRNA-seq technologies are still rapidly evolving^2^, and one of the most pressing challenges today is to address the large amount of technical noise that can drive e.g. 50% of the cell-cell variation in expression measurements^3–5^. Two such expression measurements of interest are gene detection (the identification of the set of all genes truly expressed in a given cell) and gene quantification (the estimation of the relative number of transcripts per gene and cell, also referred to as counts). The fidelity of these measurements for a given technology is termed its sensitivity and accuracy, respectively. Both sensitivity and accuracy vary widely between scRNA-seq technologies^6^, the result of the small quantities of RNA sequenced per cell compared to bulk RNA technologies, reverse transcriptase inefficiency and amplification bias^5^, among other features of the scRNA-seq protocols.

Independent of technology choice, scRNA-seq experimental design necessitates a cost trade-off between deeper sequencing of individual cells and sequencing more cells overall. We have observed that as the number of cells sequenced increases, the average gene detection rate decreases, as does the average number of molecules sequenced per cell (**Supplementary Fig. 1**), due to both choice of technology and cost trade-off. We reasoned that when the number of unique molecules drops too low, the signal-noise ratio of the data may be too low to make gene quantification informative^7^, and therefore downstream analyses should be adapted to primarily consider gene detection patterns only.

In this paper, we make the key observation that on scRNA-seq datasets exhibiting high technical noise, dimensionality reduction using only the gene detection measurements is superior to existing state-of-the-art methods that use both detection and quantification measurements^8,9^. We show that our new detection-based model, single cell Binary Factor Analysis (scBFA), leads to better cell type identification and trajectory inference, more accurate recovery of cell type-specific markers and is much faster to perform compared to quantification-based methods. Through simulation experiments, we demonstrate that our gene detection model is superior precisely when quantification noise exceeds detection noise, providing a principled explanation for when and why discarding quantification estimates is advantageous. Finally, we demonstrate the superiority of our detection model in the analysis of single cell chromatin accessibility data, suggesting detection models may improve downstream analysis of other single cell genomic modalities in high throughput datasets.

## Results

### scBFA achieves superior performance in cell type identification across diverse benchmarks

We first hypothesized that the performance of scRNA-seq analysis tools that model gene counts (quantification) could be improved by instead modeling only gene detection patterns when analyzing datasets that have a high degree of technical noise. Our intuition is that it is well established that poorly expressed genes are hard to accurately quantify using single cell genomics technologies due to technical noise^10,11^. Extrapolating to an entire dataset, we then reasoned that for datasets in which technical noise leads to low gene detection and noisy quantification, modeling differences in small gene counts is challenging and prone to error, and therefore focusing only on gene detection would be more robust.

To test our hypothesis, we developed single cell Binary Factor Analysis (scBFA), a method for dimensionality reduction that only uses gene detection patterns. We compared scBFA against seven other approaches that model gene counts and represent the spectrum of approaches to identifying cell types within scRNA-seq datasets (see Methods): scVI^12^,SAVER^13^,sctransform^14^, scrna2019^15^, PCA, ZINB-WaVE^8^ and scImpute^9^. scBFA is designed as a gene detection-based analog of ZINB-WaVE, and so comparison of scBFA versus ZINB-WaVE is the most direct comparison of gene detection versus quantification-based approaches. In this study, we focus on the task of dimensionality reduction, as it is a nearly ubiquitous first step both for data visualization and analysis^16–18^ and many analysis tools have been developed to address it^8,9,12,19,20^. Furthermore, previous work has shown that cell type identification and dimensionality reduction is still possible in scRNA-seq experimental designs favoring high cell counts, with low coverage per cell^4,21–23^.

We evaluated methods using 14 benchmark datasets for which experimentally defined cell type labels were available (**Supplementary Table 1**) by first learning low dimensional embeddings, then using the embeddings to predict cell type labels in a supervised setting. When using highly variable genes (HVG) as a gene selection criterion during data preprocessing, we found scBFA was the best, or tied for best, in 13 out of 14 benchmarks (Figs. 1-2, **Supplementary Figs. 2-3**), and this result was robust to selection of the hyperparameters of scBFA (**Supplementary Fig. 4**). Surprisingly, we found that the choice of gene selection had a significant impact on our results. Under an alternative gene selection procedure that biases towards highly expressed genes (HEG) and selects gene sets with minimal overlap with HVG (**Supplementary Table 2**), scBFA was a top performer in only nine of fourteen of the benchmarks (**Supplementary Fig. 5**).

**Figure 1:**
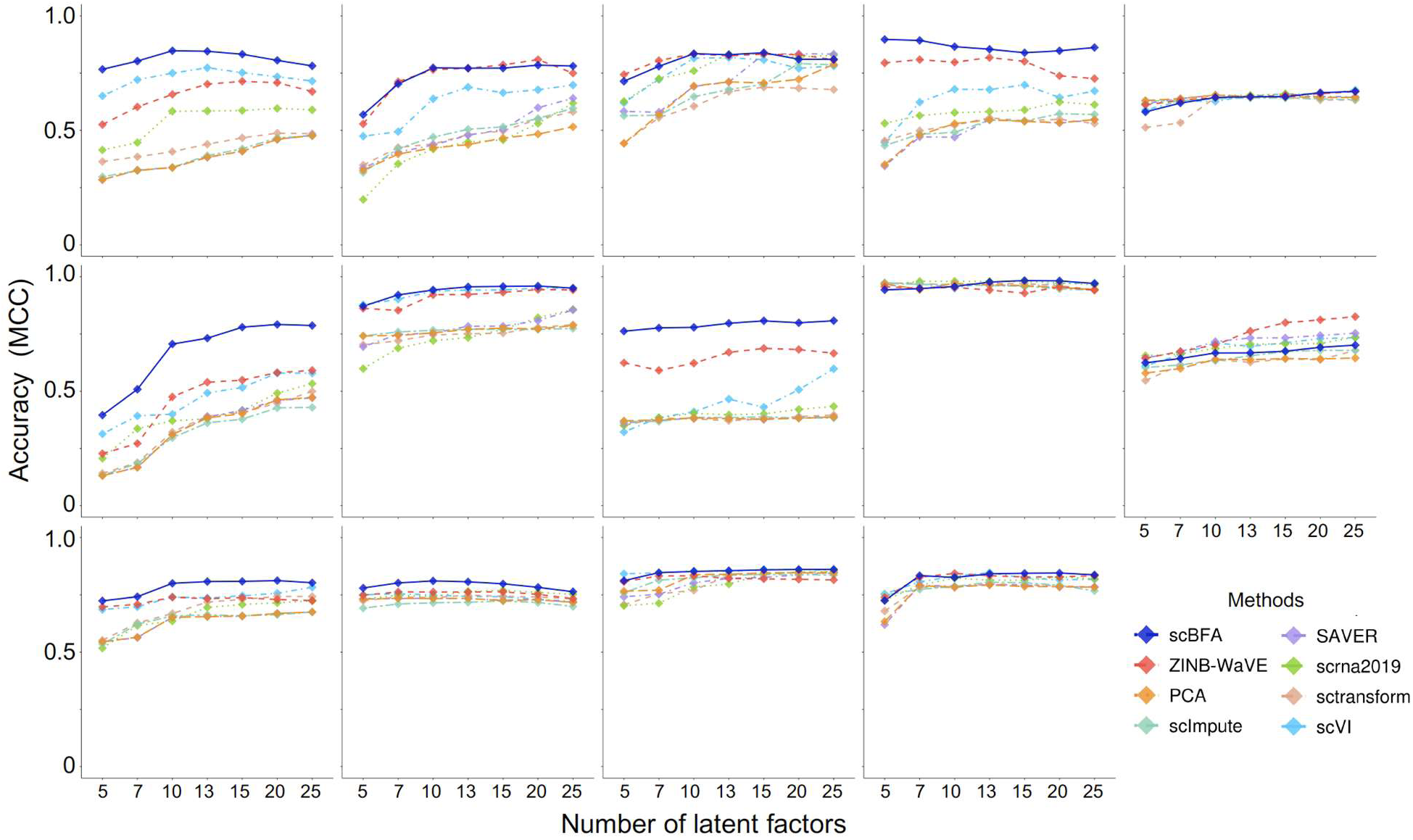
Single cell Binary Factor Analysis (scBFA) outperforms quantification models. Performance is measured via cross-validation of cell type classifiers trained on scRNA-seq benchmark data in the respective embedding spaces of each method, as a function of the number of latent dimensions specified. scBFA is top performer in 13 out of 14 datasets. Datasets from left to right, top to bottom: Dendritic, Pancreatic, DC, mESCs, HSPCs, MGE, Intestinal, MEM-T, H7-ESC, LSK, Myeloid, HSCs, PBMC, and LPS (see **Supplementary Table 1**).

**Figure 2:**
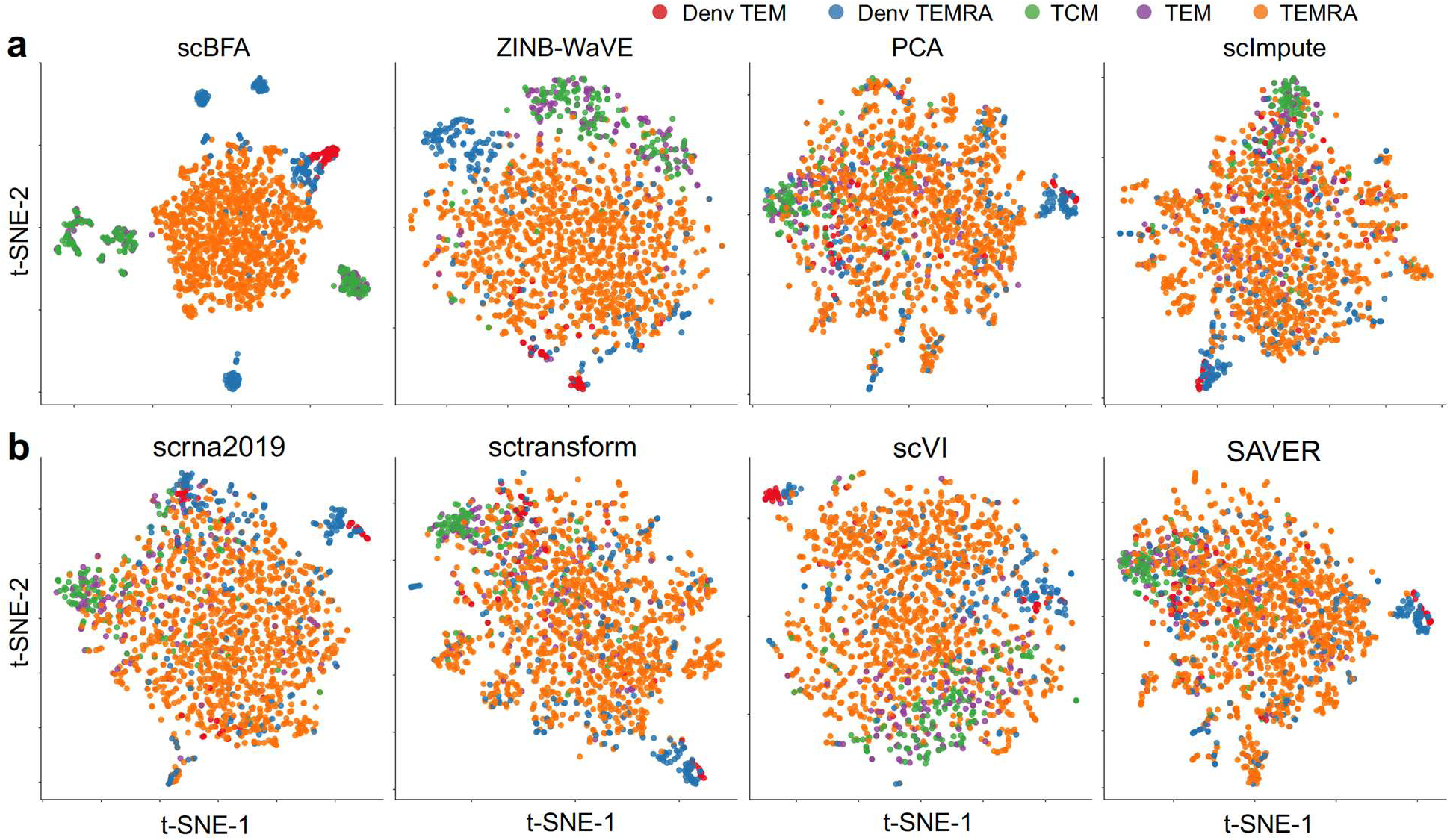
scBFA improves visualization of cell identity in the MEM-T benchmark of Patil et al. 2D t-SNE visualization of 10-dimensional embeddings generated by the eight methods on the MEM-T benchmark. Cells are colored according to their corresponding cell types and states.

### Gene selection shapes cell type identification performance by modulating detection rate and dispersion

We hypothesized the stark difference in performance between the HVG and HEG selection criteria was due to differences in overall technical noise in the resulting selected gene sets. For both the HVG and HEG versions of each benchmark, we computed two indirect measures of technical noise, the gene detection rate (GDR) and gene-wise dispersion. Existing approaches to directly estimating technical noise require spike-in standards^24,25^, and not all datasets we analyzed had incorporated spike-in standards in their protocol. GDR is the average fraction of genes that are detected as expressed in a given cell, and includes technical dropout events (genes whose expression is not detected, although they are truly expressed) as well as genes truly not expressed in a cell. Gene-wise dispersion is intuitively the excess variation in gene expression observed beyond what is expected based on a Poisson model of sampling noise, and is driven by both technical noise and biological factors of interest.

When considering each individual benchmark in isolation, HVG selection leads to a systematically lower GDR and higher gene-wise dispersion compared to HEG selection (Fig. 3). Furthermore, HEG selection consistently leads to higher performance in cell type identification for all methods tested (Fig. 3c), suggesting that HEG selection may be more sensible for cell type identification. This result is intuitive, as the HVG selection procedure identifies genes whose variance is in excess of that predicted by sampling noise, and therefore is likely to be enriched in poorly expressed genes that exhibit significant dropout noise. For scBFA specifically, HVG only outperformed HEG for the three datasets with highest GDR (Fig. 3c), consistent with our intuition that scBFA performs best when GDR is lower and therefore technical noise is higher.

**Figure 3:**
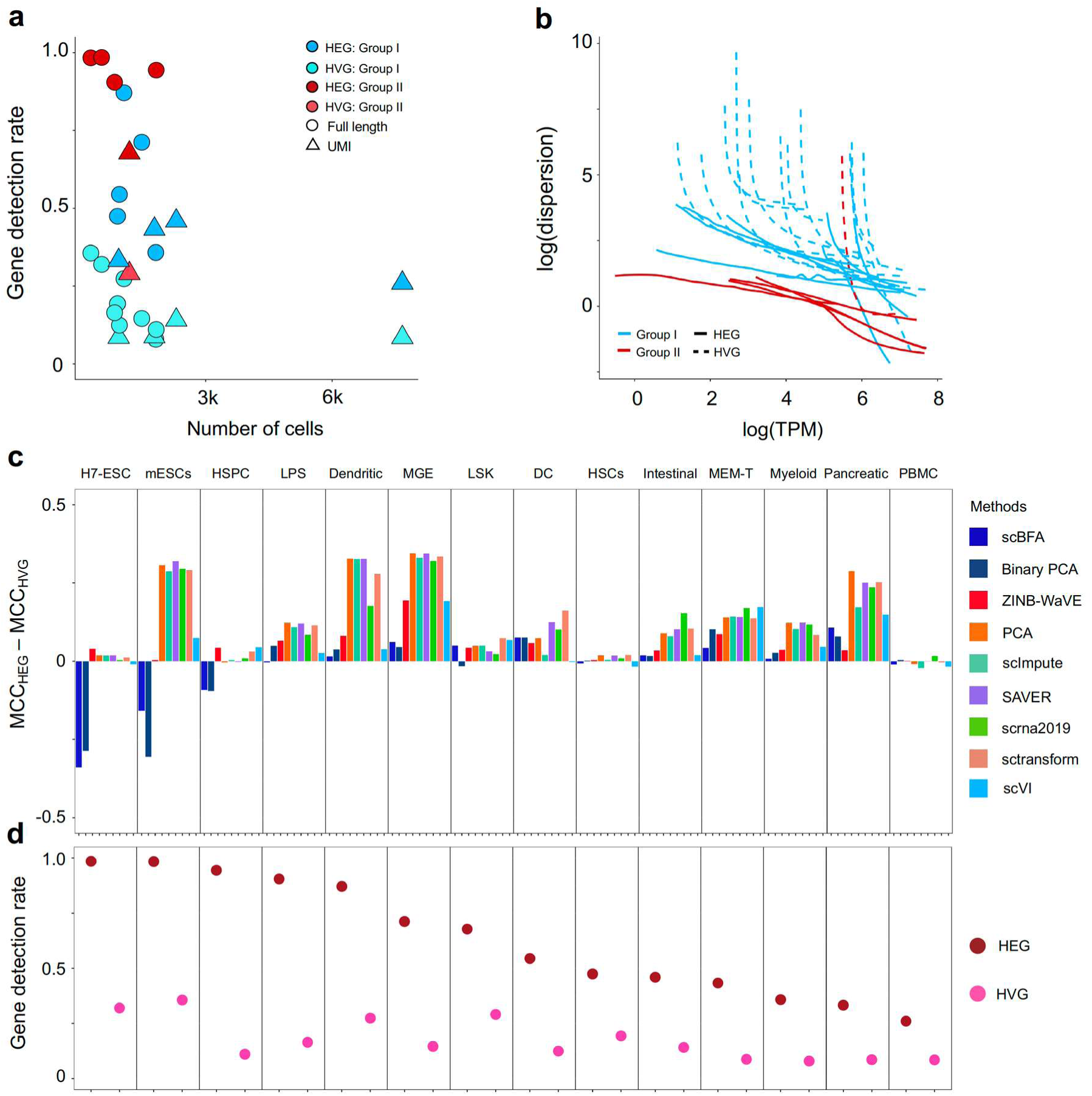
Relative scBFA performance is positively correlated with dataset size and high technical noise. **(a)** Gene detection rate as a function of the number of cells for the fourteen benchmarks, when processed using either HVG or HEG selection. Group I benchmarks refers to those datasets in which scBFA is a top performer, and Group II benchmarks refers to datasets in which scBFA is a poor performer. Note that each of the fourteen benchmarks are represented twice (once under each of HEG and HVG selection). **Supplementary Table 6** indicates the membership of each benchmark within Group I and Group II. **(b)** Same as (a), except mean-dispersion trends are estimated and visualized for each of the datasets from Group I and Group II, under HVG and HEG selection criteria. **(c)** Difference in performance (MCC) of cell type classifiers trained on individual benchmarks and for each method, either using HVG or HEG selection. Performance is assessed through cross-validation of cell type classifiers trained on scRNA-seq data in the respective embedding spaces of each method. Performance is averaged across all number of latent dimensions tested. **(d)** The corresponding gene detection rate under the two gene selection criteria. Note HEG yields systematically higher GDR compared to HVG.

Across all benchmarks, we found that scBFA outperforms count-based methods for benchmarks with low GDR and high gene-wise dispersion (Fig. 3). This is likely because higher dispersion increases the noise within gene counts, therefore forcing count-based models and their low dimensional embeddings to explain more outliers and noise in the data; this is particularly true for count models that share variance parameters across genes^8,26^. On the other hand, the gene detection pattern is more robust to noise than counts because moderate to highly expressed genes are likely to be equally well detected even in the presence of technical noise. Interestingly, low GDR of a dataset in particular is associated with more sequenced cells regardless of the experimental protocol used (**Supplementary Fig. 1**) and is likely a result of investigators trading off sequencing many cells at the cost of sequencing fewer reads per cell. These results together suggest scBFA is more appropriate for large-scale datasets.

Inversely, high GDR is more typical of smaller datasets (**Supplementary Fig. 1**) and yields poor performance of scBFA. This is because when the GDR reaches close to 100%, every gene is detected in nearly every cell, so there is limited variation for scBFA to capture in its embedding space.

### Balance of detection and quantification noise determines the relative performance of detection and count models

We next sought to identify precisely which types of technical noise were responsible for the relative performance of scBFA versus the gene count models. Previous studies found that sensitivity and accuracy (gene detection and quantification) can be affected differently by e.g. sequencing depth and other features of the protocols^7,27^. We hypothesized that differences in detection and quantification noise might explain the performance difference between scBFA and the quantification-based methods. Because technical noise is difficult to estimate in real datasets without the spike-in standards, we instead generated thousands of simulated scRNA-seq datasets that systematically vary in the relative amount of noise in gene detection and gene counts (quantification).

Our simulation framework extends the ZINB-WaVE statistical model^8^ to include parameters that separately influence the noise added to either the gene detection pattern 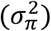 or the gene counts 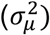 in the simulated datasets. We also tuned the global level of gene dispersion that drives variation in gene counts via the parameter *r*, which adds noise specifically to the UMI counts in the dataset and is a key parameter of many dimensionality reduction models^8,12,28^. Finally, we also tuned the global level of gene dropout observed in the dataset via the parameter *δ*, to simulate global differences in gene detection typically observed between different protocols and technologies^6^.

We first confirmed that our simulation framework generates datasets with similar characteristics to real datasets. For each of the LSK, HSPC, and LPS benchmarks, we first applied the HVG selection procedure and fit the ZINB-WaVE model. Using the ZINB-WaVE learned parameters, as well as after setting our additional framework parameters (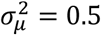, 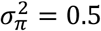, *r* = 1, *δ* = −0.5), we then simulated the exact same number of cells as was in the original dataset. Upon performing dimensionality reduction and visualization of both simulated and measured cells simultaneously, we found cells clustered by cell type regardless of whether they were from the real or simulated dataset (**Supplementary Fig. 6-8**), confirming our simulation framework generates realistic datasets.

scBFA consistently outperforms the count-based methods in classifying cell types precisely when the gene detection noise is less than the gene count noise 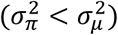 (Fig. 4). This observation is robust to the choice of gene dispersion parameter *r* (**Supplementary Figs. 9-10**) and gene selection procedure (Fig. 4, **Supplementary Figs. 11-13**). On real datasets, we found scBFA performance increases as the gene detection rate decreases (Fig. 3a), suggesting that in the real datasets for which GDR is low, the count noise may exceed the detection noise. Consistent with these results, we found that the performance of scBFA decreases after imputation (SAVER-scBFA, scImpute-scBFA) relative to before imputation (scBFA) (**Supplementary Fig. 14**), in part because imputation decreases technical variance in gene detection space.

**Figure 4:**
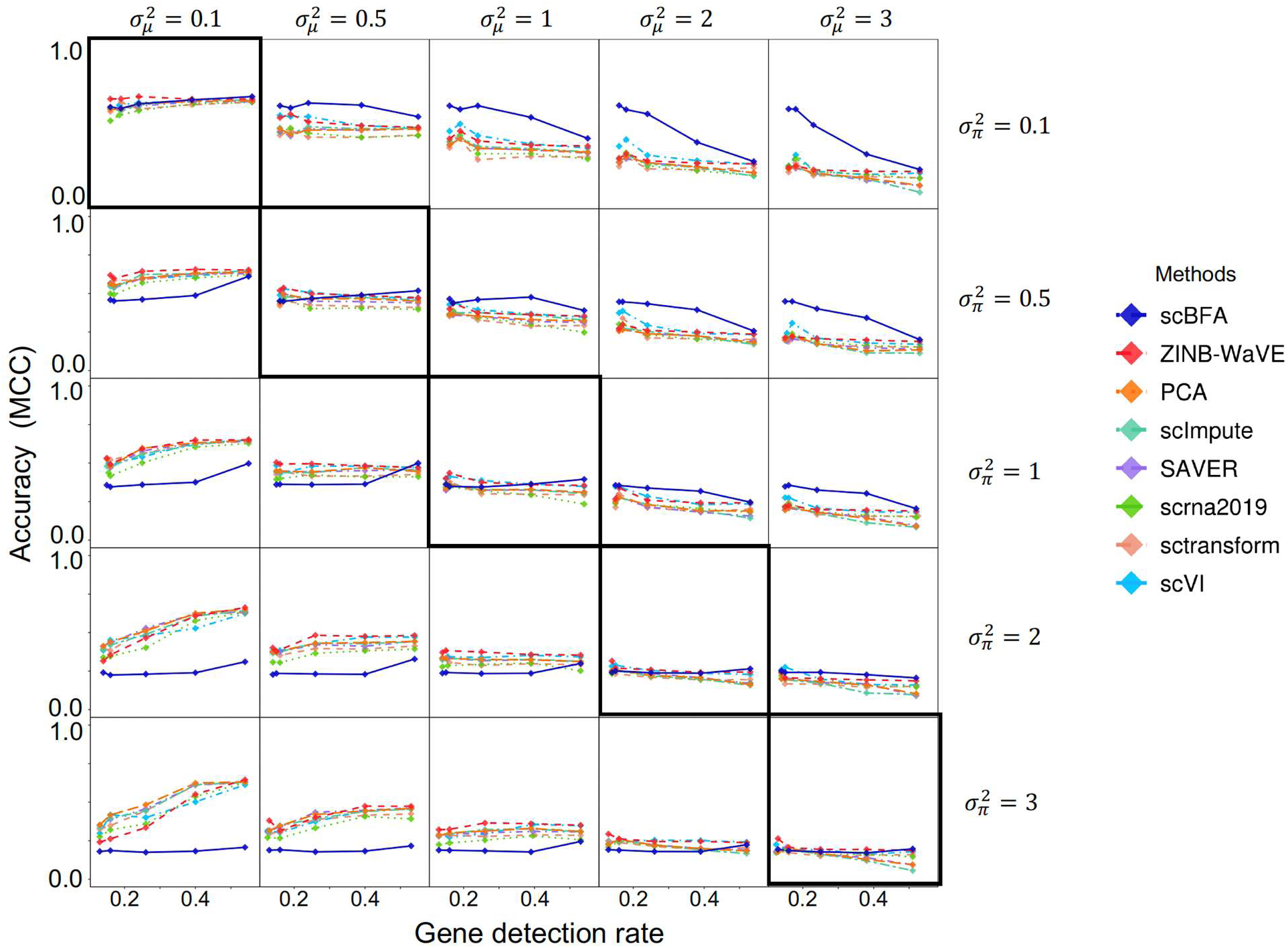
scBFA outperforms quantification models when the gene detection noise is less than gene quantification noise. Rows represent different settings of (gene) detection noise 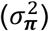, and columns represent different settings of (gene) quantification noise 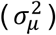. The diagonal represents simulations where the detection noise is equal to the quantification noise 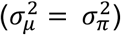, and the plots above the diagonal represent simulations where the detection noise is less than the quantification noise. The y-axis indicates the cross-validation performance of cell type predictors trained on embeddings learned from the simulated data, while the x-axis represents the gene detection rate that is manipulated by the parameter δ. Here, the ground-truth embedding matrix is obtained by fitting ZINB-WaVE to the LPS benchmark under HVG selection. The dispersion parameter r is set to be 1 in these simulations.

### scBFA mitigates technical and biological noise in noisy scRNA-seq data

We next tested each method’s ability to reduce the effect of technical variation on the learned low dimensional embeddings, by training them on an ERCC-based dataset^29^ with no variation due to biological factors. In this dataset, ERCC synthetic spike-in RNAs were diluted to a single concentration (1:10) and loaded into the 10x platform in place of biological cells during the generation of the GEMs. This dataset therefore only consists of a single “cell type”, with only technical variation present (since the spike-in RNAs were diluted to the same concentration). **Supplementary Figure 15** illustrates that both scBFA and Binary PCA yields a low-dimensional embedding with minimal variation between “cells” compared to the other methods, suggesting that gene detection models are systematically more robust to technical noise compared to count models.

We also found that modeling gene detection patterns helps to mitigate the effect of biological confounding factors in the scRNA-seq data. For example, a common data normalization step is to remove low-quality cells for which many reads map to mitochondrial genes, as these cells are suspected of undergoing apoptosis^30^. However, finding a clear threshold for discarding cells based on mitochondrial RNA content is challenging (**Supplementary Fig. 16**). We found that low dimensional embeddings learned by count-based methods are clearly influenced by mitochondrial RNA content, but this is not true for scBFA (**Supplementary Figs. 17-18**), suggesting that scBFA analysis of data will make downstream analysis more robust to inclusion of lower quality cells.

### scBFA embedding space captures cell type-specific markers

We further hypothesized that scBFA performs well at cell type classification in high quantification-noise data because detection pattern embeddings are purely driven by genes only detected in subsets of cells such as marker genes, while this is less true for count models. Marker genes should always be turned off in unrelated cell types, and always be expressed at some level in the relevant cells.

To test our hypothesis, we measured the extent to which learned factor loadings capture established cell type markers on the PBMC, HSCs and Pancreatic benchmarks, for which clear markers could be identified. For these three datasets, we identified 41, 43 and 73 markers respectively from the literature (**Supplementary Tables 3-5**). Figure 5 demonstrates that for these three datasets, the embeddings of scBFA are driven by cell type markers more than the quantification-based methods, despite the fact that the cell type markers are not used when learning the embeddings. These results also hold when HEG selection is used instead of HVG (**Supplementary Fig. 19**).

**Figure 5:**
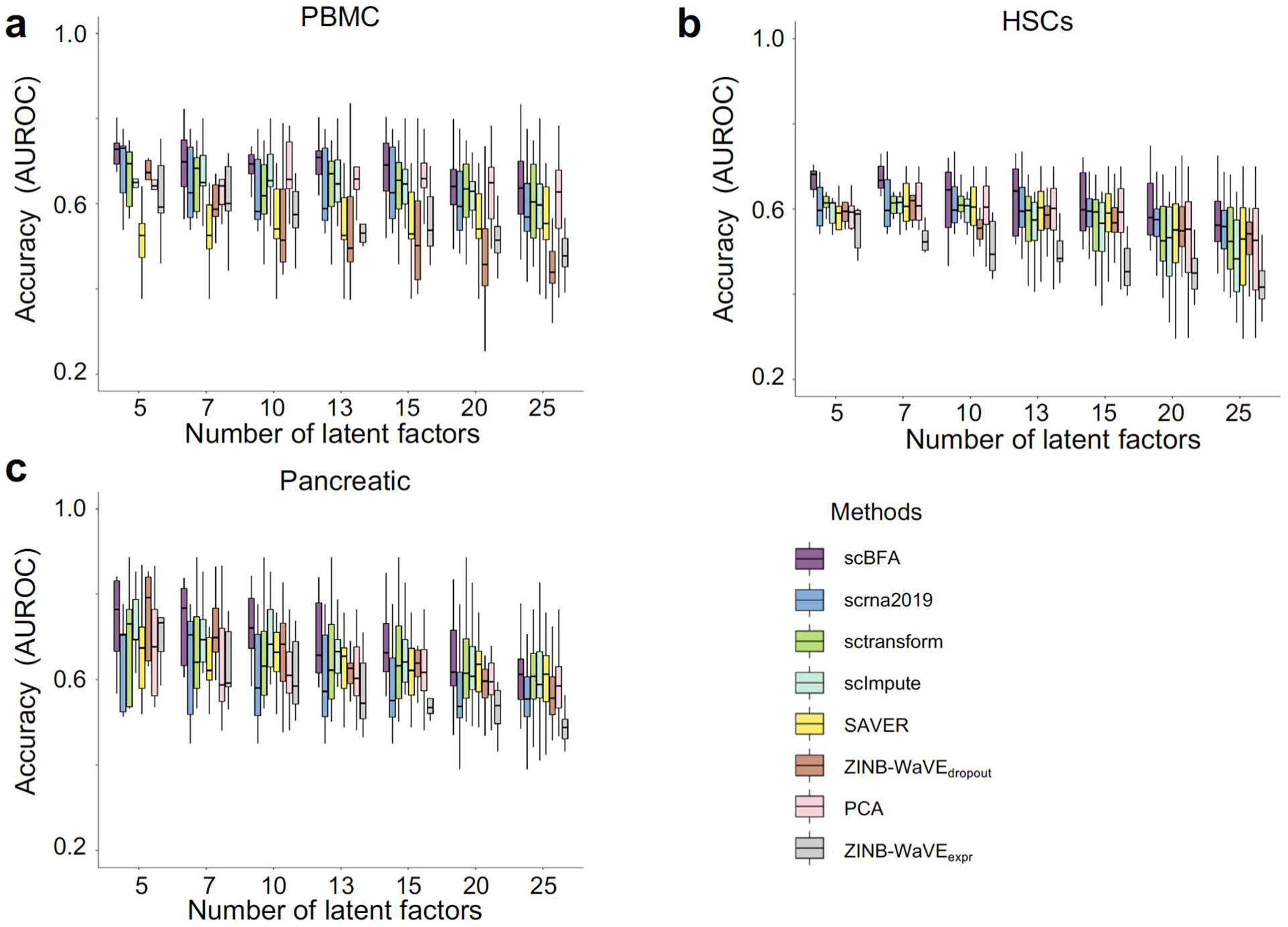
scBFA is better informed by cell type markers than quantification models. Each latent factor learned from each method was evaluated based on how much influence established cell type markers had on its embeddings, as measured by the area under the curve (AUROC) metric. Each boxplot represents the AUROC of all latent factors for a given method, for a given benchmark. ZINB-WaVE is represented twice, once for the latent dimensions inferred by their gene detection pattern (ZINB-WaVE_dropout_), and once for the latent dimensions inferred from the gene counts (ZINB-WaVE_expr_). (a) PBMC benchmark. (b) HSCs benchmark. (c) Pancreatic benchmark.

An important conceptual difference between scBFA and quantification-based methods such as ZINB-WaVE is that scBFA treats all zero-count measurements as true observations in which a specific gene is truly not expressed in a given cell. In contrast, ZINB-WaVE and others try to statistically distinguish dropout events from true zero-count measurements. As a result, the ZINB-WaVE model has a gene detection specific feature matrix and gene count specific feature matrix component, and we compared the performance of each component individually with respect to cell type marker identification. Figure 5 illustrates that scBFA factor loading matrix outperforms both components of ZINB-WaVE, suggesting the proportion of false-positive (undetected) zero-count measurements is relatively small and hard to infer statistically.

### Trajectory inference improves with detection modeling

One of the most tantalizing applications of scRNA-seq is trajectory inference for identifying changes in gene expression during continuous processes such as differentiation^31^. There are on the order of at least 45 methods for trajectory inference^32^. A first step to many trajectory inference methods is dimensionality reduction, of which PCA is a commonly used method^31^. Using a recent benchmark of trajectory inference methods, we identified a top performing method that uses dimensionality reduction (Slingshot^33^), and evaluated its performance on a set of 18 “gold standard” trajectory inference benchmarks when we replaced its PCA step with one of the dimensionality reduction methods we have benchmarked^32^. We found that substituting scBFA in place of PCA led to systematically higher performance compared to the other methods (ZINB-WaVE, PCA, scImpute, SAVER, scrna2019, sctransform, scVI) (Fig. 6). These results are robust to performance metric (Fig. 6, **Supplementary Fig. 20**).

**Figure 6:**
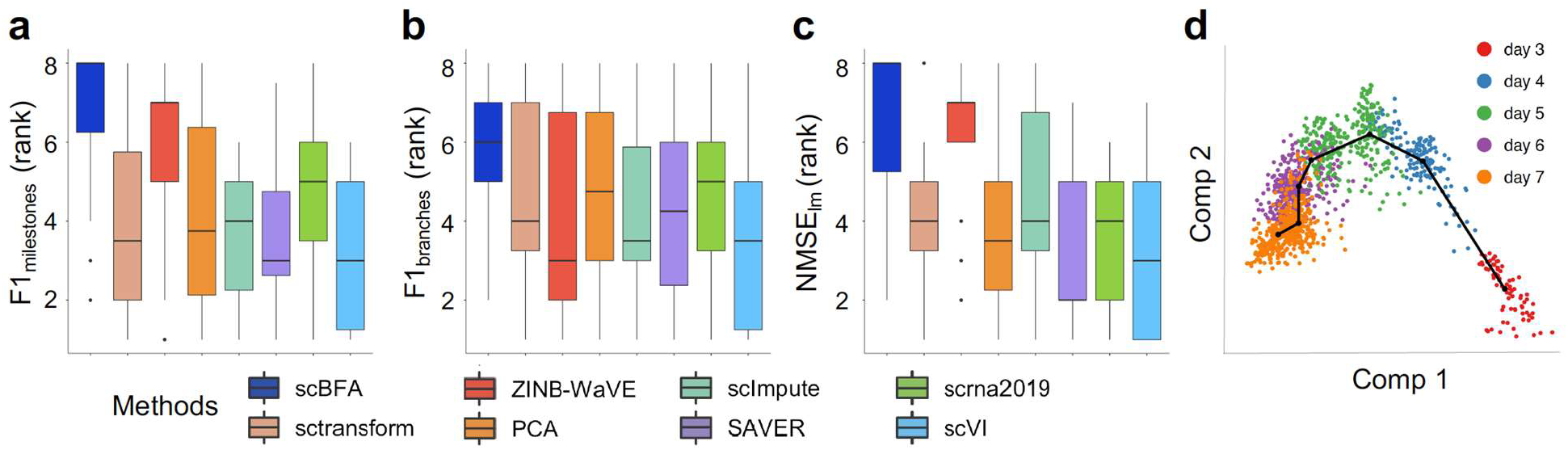
scBFA leads to the most improvement in trajectory inference performance of Slingshot. **(a-c)** Slingshot was modified by replacing the PCA step of the original Slingshot method with each of the dimensionality reduction methods tested. The y-axis shows the distribution of overall ranks (higher rank is better) of the modified versions of Slingshot. Methods were evaluated across 18 “gold standard” benchmarks and using three different performance metrics, F1_milestone_, NMSE_lm_ and F1_branch_, that measure how well the inferred trajectory matches the ground truth trajectory. F1_milestone_ and F1_branch_ are based on the quality of clustering of cells in the trajectory, while NMSE_lm_ assesses how well the position of a cell in the inferred trajectory predicts the position of the cell in the ground truth trajectory. Across the three evaluation metrics and 18 benchmarks, scBFA yields better overall performance (rank). **(d)** A 2D scatter plot of scBFA’s first two components, visualizing the inferred trajectory corresponding to the embryo development time in the H-embryos dataset.

### Detection pattern models are also superior for scATAC-seq data analysis

Several of the features of scRNA-seq protocols thought to drive technical noise are also shared amongst other single cell genomic technologies, such as small starting material and amplification bias. We therefore hypothesized that detection-based approaches such as scBFA are applicable to other types of single cell genomic data. We measured scBFA’s ability to cluster cells into cell types using scATAC-seq datasets, which also typically produce highly sparse datasets. scATAC-seq datasets are not typically suitable for input into scRNA-seq analysis tools, because the largest values observed in scATAC-seq data correspond to the ploidy of the genome (e.g. two for humans). However, such sparse, small count data means that transformation into a detection pattern matrix suitable for input into e.g. scBFA will not alter the input data significantly, making scBFA potentially more generalizable than other scRNA-seq analysis tools.

We performed dimensionality reduction and cell type classification experiments on several scATAC-seq datasets, analogous to our scRNA-seq analyses above. We benchmarked scBFA against PCA, Binary PCA, Scasat^34^, Destin^35^ and scABC^36^. scBFA systematically outperformed all other methods in our benchmark datasets (Fig. 7, **Supplementary Figs. 21-22**). An important advantage of scBFA over the other scATAC-seq methods is that only scBFA can systematically adjust for cell-level covariates such as QC measurements (e.g. cell cycle stage) and batch effects. In contrast, other methods such as Scasat are unable to remove batch effect in all features since Scasat removes batch effects through removing batch specific loci, which can be confounded with cell type-specific loci depending on the experimental design. Methods such as Binary PCA cannot directly regress out continuous covariates.

**Figure 7:**
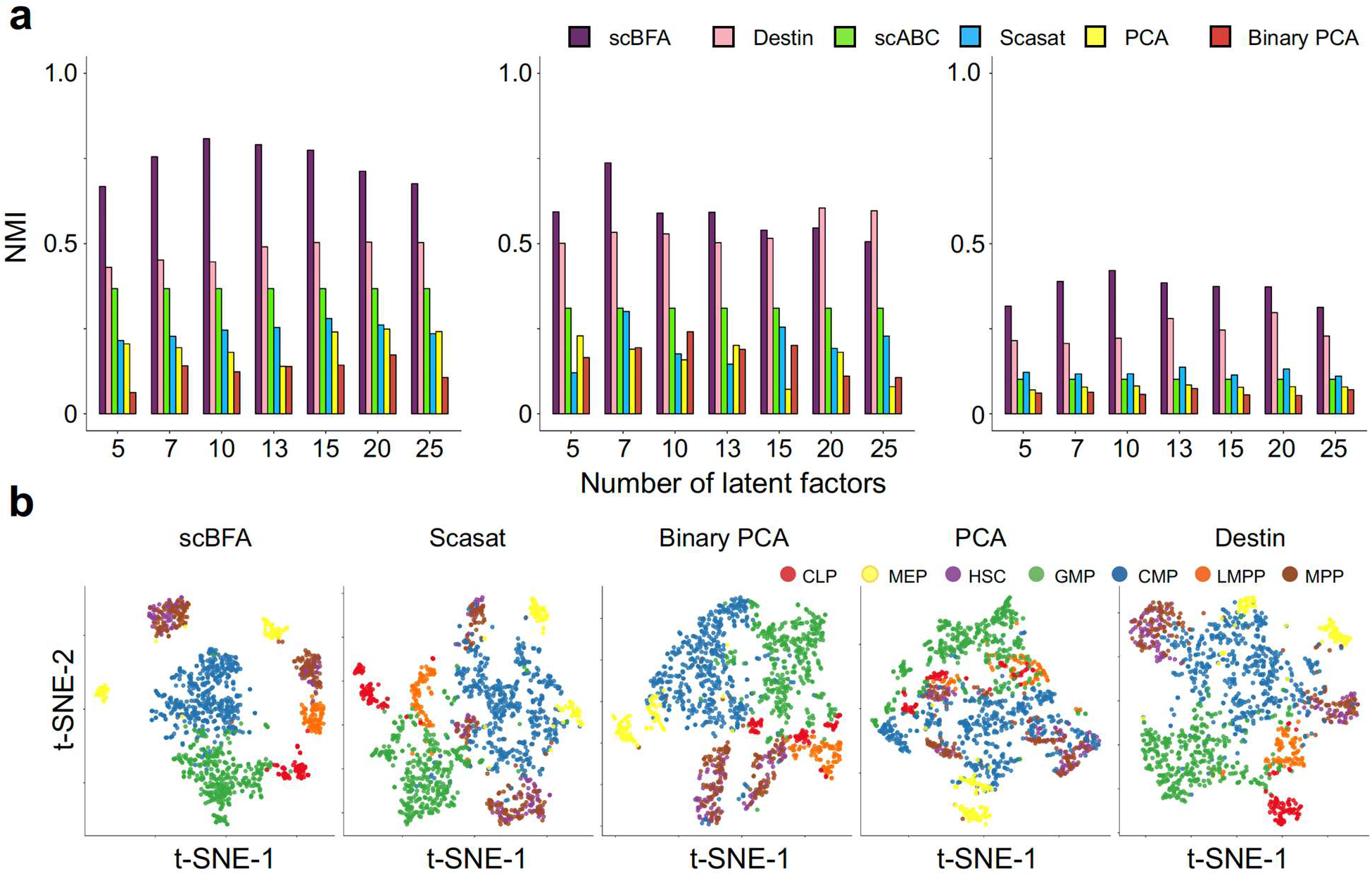
scBFA more accurately recovers cell type identity in scATAC-seq datasets. **(a)** Clustering accuracy (NMI) of each scATAC-seq method trained on the scATAC-seq dataset, as a function of the number of latent dimensions specified. **(b)** 2D t-SNE visualization of 10-dimensional embeddings generated by the five methods on dataset GSE96769. Cells are colored according to their corresponding cell types and states. scABC is omitted because it does not perform dimensionality reduction.

### Detection pattern models are faster to train by orders of magnitude

The size of scRNA-seq datasets is regularly climbing to new scales year over year^2^, as newer technologies increase the throughput of cells. With current datasets occasionally exceeding one million cells, computational efficiency of scRNA-seq analyses becomes challenging as ideally these tools could be run on local machines. We therefore benchmarked methods in terms of their speed of computation. In our comparisons, we also included a fast approximation of scBFA, which we term Binary PCA. Binary PCA is easy to implement in one line of R code (we simply transform the gene counts into gene detection patterns as a preprocessing step before use of PCA), and provides immediate benefits over standard PCA and other methods with respect to cell type identification (**Supplementary Figs. 23-24**). **Supplementary Figure 25** shows that Binary PCA is tied for the fastest of all methods, while scBFA is still faster than several competing count-based methods. More specifically, scBFA is a median of 10 times faster than ZINB-WaVE. The difference in execution time between scBFA and ZINB-WaVE is due primarily to the additional burden of modeling gene quantification because the scBFA model structure and parameter learning algorithm was designed to match the gene detection pattern component of ZINB-WaVE as closely as possible. This suggests gene detection models may help analysis tools scale to larger datasets in the future.

## Discussion

Our primary result is that when the count (quantification) noise is relatively high in a dataset as is typical in larger datasets, the effects of this noise on downstream analysis can be mitigated by modeling detection patterns instead of counts. The improvement in performance of scBFA over ZINB-WaVE in this regime (Figs. 1, 3) is particularly informative because the ZINB-WaVE model has two components: one that models gene detection, and the other that models gene counts (quantification). The model structure and parameter learning algorithm of scBFA is designed to match the gene detection component of ZINB-WaVE as closely as possible, making the difference in their performance primarily due to whether gene quantification (ZINB-WaVE) or gene detection (scBFA) is modeled.

We show that as the number of cells sequenced increases within a dataset, the technical noise in the data (as measured indirectly by the gene detection rate and gene-wise dispersion) and the relative performance of scBFA generally increase as well. Because many scRNA-seq applications benefit from higher numbers of sequenced cells, there is a steep upward trend of scRNA-seq dataset sizes, with some recent datasets containing nearly a million cells^2^. Our results therefore suggest that it is increasingly important that next-generation scRNA-seq analysis tools exploit the advantages of gene detection-only modeling in order to mitigate the effects of technical noise within the data. Also, given the influence of technology and protocol choice on technical noise in scRNA-seq data^6^, our results imply that future scRNA-seq tools could be designed to take advantage of the specific noise structure implied by different scRNA-seq protocols, as opposed to being relatively protocol-agnostic as they are today.

While it is challenging to measure technical noise in real datasets, we show that the gene detection rate and gene-wise dispersion are easily calculated and serve as good proxies for measuring technical noise. In our results, we found that the performance of scBFA exceeds that of gene count analysis tools in cell type classification when the gene detection rate falls below 90%, therefore providing the community with a specific guideline for when detection-based tools such as scBFA should be used instead of quantification-based tools. Our R package also has implemented a function, diagnose, to assist users in determining whether scBFA is appropriate for their data. Our results are also consistent with previous work that shows tasks such as dimensionality reduction, cell type identification and abundance estimation can be performed successfully when individual cells are sequenced to shallow depth^4,21–23^ and further provide a complementary analysis approach suitable for these datasets with low per-cell sequencing depth.

There is a plethora of data normalization methods that have been, and continue to be, designed to decrease technical noise within and across cells, in order to better perform both gene detection and quantification, and to make these quantities comparable across cells (see ^5,37,38^ for an overview). The challenge we address here is not addressed by data normalization methods however, as we argue that when the number of UMIs sequenced per cell decreases drastically, gene quantification information specifically is not present (or useable) in the data, which is a problem that data normalization cannot mitigate. Data normalization works complementarily to gene detection pattern analysis however, as illustrated by our use of data normalization before gene detection modeling in this work.

Our results also imply that models that statistically distinguish dropout events from genes truly not expressed in a cell may be less fruitful for large datasets. Many gene count-based methods^9,24,39^ model zero counts as a mixture of genes truly turned off (biological signal) and genes that are truly expressed but not detected due to technical artifacts from the experimental protocol (technical noise)^39^. On the contrary, gene detection pattern methods such as scBFA treat all zeroes as biological signal, a key feature motivated by the observation that zero measurements driven by technical noise tend to occur for genes that are poorly expressed anyways^10,11^. The superior performance of scBFA when the gene detection rate is low suggests that for these datasets, there is not enough information in the gene counts to reliably detect technical dropout events, and therefore traditional mixture modeling can be unhelpful for high throughput datasets where gene detection rates are low.

The success of modeling gene detection patterns in scRNA-seq is not tied to a specific model structure. The performance improvement of scBFA over ZINB-WaVE, and Binary PCA over PCA demonstrate our results hold across multiple model structures and loss functions. In both cases, not only do we observe performance gains for large scRNA-seq datasets, but there is also a substantial speed improvement because detection modeling avoids complex parametric modeling of gene counts, making detection models scalable to larger datasets. Within the class of gene detection-based models, Binary PCA provides a moderately accurate but much faster and simpler implementation scheme that can be achieved in one line of code, making our results readily achievable by current analysis pipelines.

A surprising finding was that HEG gene selection led to systematically better cell type identification for every tested method in almost all datasets, compared to HVG selection (Fig. 3c). HVG selection anecdotally is the standard criterion upon which variable genes are typically selected during preprocessing^40^ suggesting at least for cell type identification, HEG selection may lead to improved performance regardless of the method used.

While single cell genomic data from different modalities such as scATAC-seq have similar data structure as scRNA-seq data, the analysis tools and pipelines developed to date for those two technologies are largely mutually exclusive. Here we show that scBFA generalizes to other single cell genomic modalities and outperforms existing methods for cell type identification for scATAC-seq datasets as well, even those that take advantage of auxiliary data such as distance to transcription start sites^35^. We expect our results to generalize to other single cell genomic modalities such as single-cell methylation or histone modification data.

## Methods

### Single cell Binary Factor Analysis (scBFA) model

scBFA is available as an R package from GitHub (https://github.com/quon-titative-biology/scBFA), and is currently under submission as a Bioconductor package. The main function to run scBFA is scbfa().

In our notation below, matrices are represented by upper case bold letters, vectors by lower case bold letters, and numeric constants as upper case non-bold letters. Square brackets also indicate a matrix, though represented as a series of column vectors. A matrix subscript with round brackets indicates the index of the corresponding column vector.

The schematic of our single cell Binary Factor Analysis (scBFA) model is shown in **Supplementary Figure 26**. The input data to scBFA consists of two matrices, ***O*** and ***X***. ***O*** is a matrix of counts, consisting of *G* features (genes in the case of scRNA-seq data, or loci in the case of scATAC-seq data) measured in each of *N* samples (cells). From the input data ***O***, we compute a matrix ***B***, where *B*_*ij*_ represents the detection pattern observed for cell *i*(*i* = 1, …, *N*) and feature *j* (*j* = 1, …, *G*). For scRNA-seq inputs, *B*_*ij*_ = 1 when *O*_*ij*_ ≥ 1, otherwise *B*_*ij*_ = 0. Therefore, *B*_*ij*_ = 1 indicates that at least one read (or UMI) maps to gene *j* in cell *i* and therefore suggests gene expression. Similarly, for scATAC-seq input data, *B*_*ij*_ = 1 when *O*_*ij*_ ≥ 1, in other words, when at least one read maps to locus *j* in cell *i* (and therefore suggests locus accessibility), otherwise *B*_*ij*_ = 0. scBFA is adapted from a generalized linear model framework and is therefore capable of adjusting for batch effects and other nuisance cell level covariates. Input ***X*** = [***x***_**1**_, ***x***_**2**_, …, ***x***_***N***_]^*T*^ is a *N* × *C* cell-level covariate matrix that enables correction for *C* observed nuisance factors such as batch effects or other cell-specific quality control measurements. If there are no such cell level covariates that need to be adjusted for, ***X*** is the null matrix by default.

The intuition behind scBFA is that it performs dimensionality reduction to explain the high-dimensional detection pattern matrix ***B*** by estimating two lower-dimensional matrices: a *N* × *K* embedding matrix ***Z*** = [***z***_**1**_, ***z***_**2**_, …, ***z***_***N***_]^***T***^, and a *K* × G loading matrix ***A*** = [*a*_**1**_, ***a***_**2**_, …, ***a***_***G***_]. Here, *K* is the number of latent dimensions used to approximate ***B***_***ij***_, where *K* ≪ *G*. *u*_*i*_ and *v*_*j*_ represent the *i*^*th*^ cell level intercept and *j*^*th*^ feature specific intercept, respectively. ***u*** is therefore a vector of length *N*, and ***v*** is a vector of length G. For scRNA-seq for example, we expect ***u*** and ***v*** will implicitly model the variation of gene expression caused by library size. *μ*_*ij*_ is the mean of the Bernoulli distribution governing whether feature *j* is detected in cell *i* or not.

Formally, scBFA is defined by the following model:

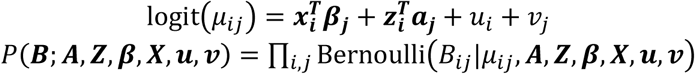

We train the scBFA model by optimizing the following penalized likelihood function:

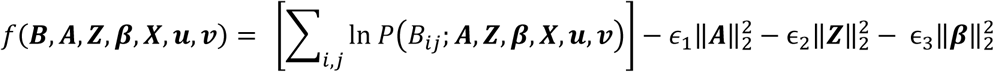

Here, ϵ_1_, ϵ_2_ and ϵ_3_ are tunable parameters that control the regularization of the model parameters, where by default 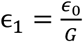, 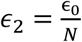, 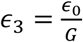, and ϵ_0_ = *max*{*N*, *G*}. The optimization is carried out using the L-BFGS-B optimization routine available in the R optim() function. After completing optimization, we orthogonalize ***Z*** and ***A*** using the orthogonalizeTraceNorm() function available in the ZINB-WaVE^8^ package.

### Binary PCA model and calculation of the gene detection pattern matrix

Binary PCA describes our fast approximation to scBFA by simply running PCA, with the exception of converting the input count matrix into a detection matrix by converting non-zero values to one. We implemented Binary PCA through the addition of a single line of R code. Suppose that countMatrix is the name of the matrix in R that stores e.g. the UMI counts for each gene in each cell. To run Binary PCA, we first convert the countMatrix into the gene detection pattern matrix before running PCA via the R command:

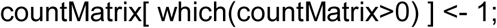

We then call PCA using the following command in R:

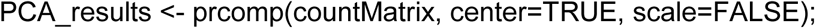

Note that typically scale is set to TRUE when calling PCA. For Binary PCA, we set it to FALSE because variance in gene detection is potentially associated with cell types (e.g. genes with higher detection variance are more likely to be marker genes, and therefore should contribute more to the embedding).

### Execution of scRNA-seq analysis methods

We compared scBFA against scVI^12^, SAVER^13^, sctransform^14^, scrna2019^15^, PCA, ZINB-WaVE^8^ and scImpute^9^. These seven methods were selected to represent diverse classes of approaches to scRNA-seq data analysis, including dimensionality reduction methods (PCA, ZINB-WaVE, scVI), preprocessing approaches that can be applied before dimensionality reduction (sctransform, scrna2019), and imputation methods that can be applied before dimensionality reduction (SAVER, scImpute). Of the dimensionality reduction methods, PCA was chosen because of its implementation in popular packages such as Seurat^40^, and scVI^12^ is a leading deep learning-based dimensionality reduction method. ZINB-WaVE was chosen specifically because it is a recently-developed method and scBFA is designed as a gene detection-based analog of ZINB-WaVE, and so comparison of scBFA versus ZINB-WaVE is the most fair comparison of gene detection versus quantification based approaches.

We ran most of the scRNA-seq analysis methods with their default parameter settings, with the exception of scVI and scrna2019.

scVI requires specification of a learning rate and the number of iterations before convergence. During training, we found scVI performance was heavily influenced by these two parameters. We therefore performed an unbiased grid search by setting the number of iterations to be either 2000 or 4000, and seting the learning rate to be either 1e-2, 1e-3 or 1e-4. We then trained the model with all six possible combinations of learning rate and number of iterations, and selected the combination of parameters that gave the lowest loss on the hold-out set. The loss value is provided by scVI during training. During training, the size of the training set is fixed to be 75% of the entire dataset, and the remaining parameters are fixed at their default values. We repeated the above parameter search for the same number of factors as the other methods for all scRNA-seq datasets. For the simulated datasets, given the large number of scenarios tested, we fixed the learning rate to be 0.001 and number of iterations to be 2000.

scrna2019 is a method developed to perform feature selection and GLM-based factor analysis on scRNA-seq^15^. The scrna2019 R package (obtained on May 6, 2019 from https://github.com/willtownes/scrna2019) offers both a GLM-factor analysis model and a corresponding deviance score approximation. We used the deviance score approximation instead of the GLM framework for our experiments because several benchmarks required batch effect correction, which should be addressed using the deviance score approximation as per scrna2019’s authors’ recommendations^15^. Also, at the time of writing of this paper, the GLM implementation produced errors for three of our datasets that prevented us from completing our experiments.

### Execution of scRNA-seq analysis methods

We compared scBFA against PCA, Binary PCA, Scasat^34^, Destin^35^ and scABC^36^, Scasat and Destin are scATAC-seq analysis tools primarily designed to identify cell types and differential accessibility analysis. Both methods treat dimensionality reduction as prior step before further clustering distinct cell types. Scasat’s embedding space is learned by performing multidimensional scaling (MDS) on a cell-cell Jaccard similarity matrix computed from a binarized chromatin accessibility matrix. Destin developed a weighted principle component analysis approach using distance to transcription start sites and reference regulomic data. scABC is an unsupervised clustering tool of single-cell epigenetic data and performs multi-stage clustering based on the input chromatin accessibility matrix directly. Except for PCA and scABC, all other methods binarize chromatin accessibility data in advance.

### Quantifying the effect of imputation on scBFA

We compared scBFA’s performance before and after imputation on our 14 benchmark datasets under HVG selection. We tested two state-of-art imputation methods, SAVER^13^ and scImpute^9^. SAVER estimates library size-normalized posterior means of gene expression levels 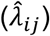, which are inappropriate for input into scBFA because they are not sparse. We therefore sampled counts from SAVER’s generative model as follows:

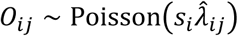

Where 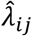 is SAVER’s imputed expression level and *s*_*i*_ is the library size for cell *i* divided by the mean library size across cells^13^. We generated five separate count matrices *O*_*ij*_ based on the SAVER estimates 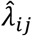. For scImpute, we used its imputed gene counts matrix directly as input for scBFA.

### Selection of representative datasets to measure gene detection rates

We obtained a total of 36 scRNA-seq datasets from which we calculated gene detection rates as a function of the number of cells in each dataset (**Supplementary Fig. 1**). We obtained these datasets from two sources, the conquer database^41^ and the Gene Expression Omnibus^42^ (GEO). For GEO, we used the search term “((‘single cell rna-seq’ OR ‘single cell transcriptomic’ OR ‘10X’ OR ‘single cell transcriptome’) AND Expression profiling by high throughput sequencing[DataSet Type]) AND (Homo sapiens[Organism] OR Mus musculus[Organism])”, sorted all datasets by size, then selected a similar number of datasets from both the top and bottom of the list (**Supplementary Table 7**).

### Computing mean and dispersion curves

We use the DEseq2^28^ package to obtain gene-specific dispersion estimates for each dataset, where dispersion is measured across all cells in a dataset. Within the DEseq2 pipeline, gene-wise dispersions are first estimated, a trend line is fit to the gene-wise dispersion estimates, and finally shrinkage is applied to the gene-wise dispersion estimates (MAP estimates). In Figure 3b, we extracted the fitted gene-wise dispersion estimates from the trend line (second step), and we fit these dispersion estimates by local linear regression (LOESS) using the gene-wise mean of transcripts per million (TPM) across all cells as the explanatory variable. To address the border effect of LOESS, we removed the top and bottom 2.5% of genes as ranked by TPM. Note that using the MAP dispersion estimates (final DEseq2 step) or the fitted dispersion estimates from the trended fit (second step) does not materially change our conclusion. The exception is for the dataset PBMC where there are 455 genes with its MAP dispersion estimates staying at their initialized value of 1e-8 during optimization, while their fitted dispersion estimates are substantially different. We therefore chose the fitted dispersion estimates to generate Figure 3b.

### Benchmarking dimensionality reduction methods for scRNA-seq

We evaluated each dimensionality reduction method by how well their low dimensional embeddings discriminate experimentally-defined cell types. For each dataset and method tested, we first performed dimensionality reduction on the entire dataset to obtain an embedding matrix representing each cell in *K* dimensions (the matrix ***Z*** described in the scBFA methods section). We then performed 5-fold cross validation in which we trained a non-regularized multi-level logistic classifier on the training embeddings from each method using the a priori known cell type labels, then used the model to predict cell type labels for the test embeddings. For every prediction, using the known cell type labels, we computed a confusion matrix and the corresponding Matthews’ correlation coefficient (MCC) as a measure of classification accuracy. MCC was calculated using the R package mltools. We repeated 5-fold cross validation a total of 15 times, and reported the mean classification accuracy as the final accuracy.

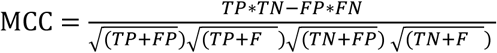

In our analysis of the ERCC dataset, we used a different evaluation metric because each “cell” represents technical replicates of the spiked-in RNA diluted at a constant ratio (10:1). Under the assumption that the only variation between “cells” is due to technical factors, we therefore used averaged within-group sum of squares (AWSS) to measure how the low-dimensional embedding learned by each method captured such homogeneity. Given a *N* by *K* embedding matrix *Z*, AWSS was calculated as follows:

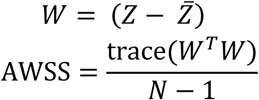

Here, 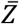 is a *N* by *K* matrix for which every row is the column mean of *Z*.

### Benchmarking cell type identification methods for scATAC-seq

We benchmarked scBFA against existing scATAC-seq analysis tools by evaluating their ability to correctly cluster cell types. We used a different evaluation scheme from that used for the scRNA-seq experiments because one of the existing methods (scABC) does not produce low dimensional embeddings, and instead outputs cluster labels. The methods Scasat and Destin both provide cluster labels directly from their analysis pipeline. For scBFA, PCA and Binary PCA, we clustered cells based on the learned embedding matrices using R’s built-in hierarchical clustering function hclust with Wald’s distance. We compared the accuracy of the clustering results from each method using the metrics Normalized Mutual Information (NMI) and Adjusted Rand Index (ARI), computed using the R package aricode.

### Simulation of scRNA-seq data

A variation of the ZINB-WaVE model was used to simulate scRNA-seq datasets and is defined as follows (**Supplementary Fig. 27**):

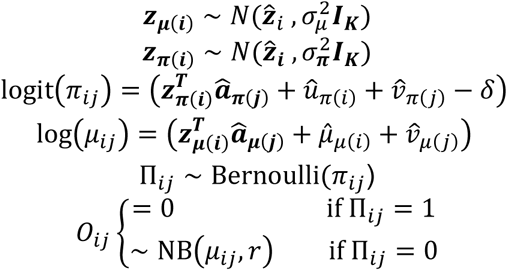

To keep the consistency of the notation, the parameters we used above 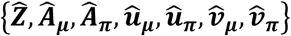 correspond to the parameters 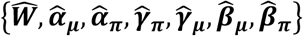 used in the original ZINB-WaVE paper, respectively. In the first step of our simulations, all parameters with a hat accent are set *a priori* by fitting the ZINB-WaVE model using its R package^8^ on a single scRNA-seq dataset in order to use realistic parameters for our simulation. The remaining parameters 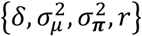 are then systematically varied in our simulations to determine their effect on downstream dimensionality reduction methods. *O*_*ij*_ denotes the gene counts for cell *i* and feature *j*. As is described in the original ZINB-WaVE paper, 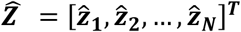 is a *N* × *K* embedding matrix, while 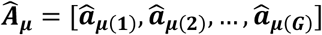 and 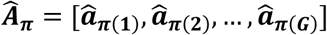 are the corresponding *K* × *G* regression coefficient matrices for the negative binomial and Bernoulli distributions governing the gene count and detection components, respectively. The output of the Bernoulli distribution is the latent variable Π_*ij*_, which decides whether a gene is detected (in which case the observed value *O*_*ij*_ is sampled from a negative binomial distribution), or not detected. 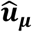 and 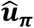 are *N* × 1 cell-specific intercepts for the count matrix and detection matrix respectively. Similarly, 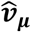 and 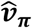 are *G* × 1 gene-specific intercepts for the count matrix and detection matrix, respectively. The number of latent dimensions *K* used to generate the gene expression values was fixed at 5, and we used a total of 2,000 highly variable genes as in the original dataset. The LPS dataset does not provide any cell level covariates, so in these simulations, there is no cell-or gene-wise covariate matrices. For quality control purposes, we filtered out genes that are expressed in fewer than 1% of the cells, and then filtered out cells in which less than 1% of genes are expressed.

The distinction between our simulation framework and ZINB-WaVE is that ZINB-WaVE maintains the same cell embedding space 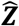 across both the gene detection and count spaces. In contrast, our framework relaxes this constraint by introducing individual embeddings ***Z***_π_ and ***Z***_μ_ that are close to 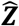. Formally, ***Z***_π_ and ***Z***_μ_ are *N* × *K* embedding matrices for the gene detection and count spaces, respectively. Each row *i* of ***Z***_π_ and ***Z***_π_ are sampled from respective *K*-multivariate Gaussian distributions with the same mean defined by 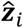 and spherical variance parameters 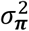 and 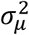, respectively.

In our simulations, we varied the simulation parameters 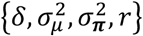 as follows. To influence the total number of gene counts detected (total detection rate), we set *δ* ∈ {−2, −0.5, 1, 2.5, 4}. To influence the variance in the gene detection and count embedding spaces, we set 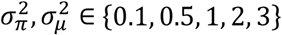. Finally, we varied the common gene dispersion parameter *r* ∈ {0.5,1,5}. In total, the number of unique parameter settings we used to simulate scRNA-seq data is 5 x 5 x 5 x 3 = 375. For each of those scenarios, we simulated 3 replicates, resulted in 375 x 3 = 1125 datasets.

### Quality control of scRNA-seq data

For each scRNA-seq dataset tested, we performed a standardized quality control process. We first removed cells for which mitochondrial genes accounted for over 50% of the total observed counts. Then we filtered out genes that are expressed in fewer than 1% of cells, and removed cells whose library size (total read or UMI count) was less than one eighth quantile of all cell library sizes. One exception is the MEM-T cell dataset, where we removed an extra 361 cells from the batch labeled “subject16” to remove batches that were confounded with cell types.

### Preprocessing of scATAC-seq data

We followed the scATAC-seq pipeline for processing and aligning reads used by the Destin method^35^, obtained from GitHub at https://github.com/urrutiag/destin on April 22, 2019. This preprocessing pipeline yielded 2779, 576, and 960 BAM files for GSE96769, GSE74310 and GSE107816, respectively. These BAM files form the initial input of Destin and scABC.

For GSE96769, we only kept cells and genomic loci that are used in the original paper’s analysis. The indices for genome loci and cells that passed quality control are supplied in the supplementary files of the original paper. Beyond that, we selected a subset of frozen cells from 5 patients, excluding patient BM0106 and a subset of pDC cells from patient BM1137 to keep as many samples as possible while removing the part of the batches confounded with cell types. This enabled us to construct a design matrix to correct for patient specific effects. We furthermore excluded cells that are labeled as unknown by the original author.

For each scATAC-seq dataset tested, we only kept genomic loci that are accessible in at least 1% of cells, and then removed cells with total number of accessible sites that deviates more than 3 standard errors to the mean (in either direction) across all cells. The number of retained cells used as input in our downstream analysis was for 1358 GSE96769, 572 for GSE74310, and 929 for GSE108716.

### Defining cell type labels in benchmark datasets

Most benchmark datasets used in our analyses were selected because the cell types were already defined in the original study by either known experimental condition or via cell surface markers. However, for the PBMC dataset, Stoeckius et al.^43^ collected single-cell antibody-derived tag (ADTs) data as well as scRNA-seq using CITE-seq^43^. ADTs can be viewed as a digital readout of cell surface protein abundance. We defined the cell types within this dataset by performing Louvain clustering on the Jaccard similarity matrix constructed based on the normalized ADTs levels, similar to Stoeckius et al^43^. Louvain clustering was performed using the “cluster_louvain” function implemented in the igraph R package. Clustering identified 10 cell types automatically. Note that the quality control standard for this dataset is different compared to the other scRNA-seq datasets used in our analysis, as cells were required to pass both scRNA-seq-specific filters (minimum of 800 reads) and ADT-specific filters (minimum of 50 ADT counts).

### Normalization of scRNA-seq data

For each method, we also normalized cells to control for differences in library size. For PCA, we normalize the counts by setting 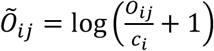, where 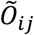 is the normalized gene count for cell *i* and gene *j*, *O*_*ij*_ is the original gene count for cell *i* and gene *j*, and *c*_*i*_ = ∑_*j*_ *O*_*ij*_ is library size for cell *i*. ZINB-WaVE directly accounts for library size via their cell-specific intercept. For scImpute, we used the total number of imputed counts per cell as their corresponding library size and normalized in the same way as PCA. For scBFA, we estimated the feature-specific intercepts and cell-specific intercepts to implicitly model the effect of library size. SAVER uses the library size divided by the median library size across all cells to adjust for cell size. sctransform uses the log of the library size in its model. scrna2019 outputs a transformed deviance score matrix that does not depend on library size as input.

### Normalization of scATAC-seq data

For PCA, we performed a log transformation 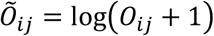 to adjust the counts within scATAC-seq, where *O*_*ij*_ is the original read count for cell *i* and locus *j*. For scBFA, Scasat, Destin, scABC, and Binary PCA, no extra normalization was applied.

### Gene selection in scRNA-seq data

Highly variable genes (HVG) selection was performed to identify the most overdispersed genes, that is, genes that exhibit more variance than expected based on their mean. The HVG selection was performed using the FindVariableFeatures command implemented in Seurat 3.0. By default, Seurat selected the top 2,000 genes. Highly expressed gene (HEG) selection was performed to identify the genes that exhibit the highest variance across cells, regardless of their mean, and is therefore expected to capture genes with higher mean expression compared to HVGs. To identify HEGs we calculated the gene-specific variance in the gene count space and select the top 2,000 genes to make the set size comparable to HVGs.

The gene detection rate (the average fraction of cells in which a gene is detected as expressed) and gene wise dispersion of each dataset calculated in Figure 3b is based on these 2,000 most variant genes under both the HVG and HEG selection criteria. For the timing experiment, we only selected the top 1,000 genes under the HVG criterion for computational speed.

### Batch effect correction

For both scRNA-seq and scATAC-seq datasets, we performed two types of batch effect correction, depending on how the cell types are distributed across the batches in the dataset. For datasets where all cell types are represented in all batches (e.g. replicates, patients), such as the HSC dataset, we used those cell level covariates to define the *N* × *C* design matrix ***X*** (see the scBFA model details above). For ZINB-WaVE, scBFA and scVI, we regressed ***X*** out within the model structure. Since PCA does not offer a framework to regress out nuisance factors, we first regressed ***X*** directly from the normalized counts 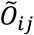 using a linear model. We then applied PCA on the residual matrix and obtained the corresponding embeddings and factor loading matrix. For Binary PCA, scImpute, SAVER, sctransform and scrna2019, we also regressed out ***X*** from the binary entries and imputed values respectively, then used the residual matrix in the same way as for PCA.

For other datasets (MEM-T, Pancreatic, MGE in scRNA-seq, and GSE96769 and GSE74310 in scATAC-seq), some batches were missing a subset of cell types, resulting in a design matrix ***X*** that cannot be directly used to estimate all batch effects. In this scenario, our strategy for modifying the dataset to address batch effects is as follows. Note that we use the same parametrization used to define the scBFA model earlier, except that we define a new observation matrix ***M*** as a *N* × *G* matrix, where observations can either correspond to measured expressed levels ***O***, inferred binary detection pattern ***B***, or imputed read counts. Except for minor differences in parameterization, the GLM-based dimensionality reduction methods scBFA and ZINB-WaVE can be summarized in the following framework, where *g* is the link function, *P* is a probability measure and *μ* is the expectation over the probability measure. In the case of ZINB-WaVE, *P* is a zero-inflated negative binomial distribution. In the case of scBFA, *P* is a Bernoulli distribution.

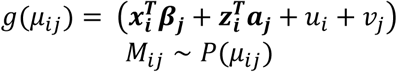

We first identify the largest subset of cell types that are represented in all batches within a given dataset. Define ***M***_sub_ as the submatrix of *N*′ observations (*N*′ < *N*) corresponding to this subset of cell types, and similarly define the submatrices ***X***_sub_, ***Z***_sub_, ***A***_sub_ and ***u***_sub_, where *i*′ = 1, …, *N*′. We ran each dimensionality reduction method once to obtain an estimate of 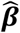 by optimizing the likelihood of the following model:

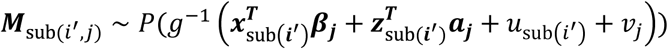

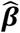 learns the variance induced by different batches only. Then, we use 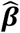 as plug-in estimate of ***β***, and performed each dimensionality method on the full dataset to obtain estimates of all other parameters. Note since both ZINB-WaVE and scBFA regularize their coefficient matrix ***β***, ***X***_sub_ and ***X*** are both standardized. For PCA, scImpute, SAVER, sctransform and scrna2019, we used a similar strategy to obtain 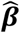 by using linear regression to regress out ***X***_sub_ from the observation matrix corresponding to the largest subset of cell types represented in all batches. Then, we calculated the residuals *R*_*ij*_ = *M*_*ij*_ − ***XB***^***T***^ on the full dataset with 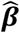 fixed and performed PCA on the residual matrix ***R***. For scVI, we were unable to modify the model framework to adjust for batch effects when they were confounded with cell types, as was the case in MEM-T, Pancreatic and MGE. Therefore, we measured scVI performance when we did not correct for batch effect, as well as when we performed naïve batch effect correction ignoring the confounding, and then reported the best performance for scVI.

Scasat handles batch effects through removal of batch-specific loci. However, for datasets GSE96769 and GSE74310, the batch effect is confounded with cell types. Therefore, we ran Scasat without batch effect correction because batch-specific loci would be indistinguishable from cell type-specific loci. Destin and scABC cannot adjust batch effect on their own, and so we ran them without batch effect correction on these two datasets.

### Identification of marker genes

We evaluated the extent to which the inferred dimensions for each method recovers known marker genes (Fig. 5). For each method, we first obtained the *K* × *G* factor loading matrix indicating which genes are contributing to each of the *K* factors. Then, for every loading matrix and given number of factors, we ranked the absolute value of each gene in each factor and calculated the AUROC (area under the receiver-operator curve) to measure the extent to which the known marker genes contribute more to a factor than expected by chance.

Note that ZINB-WaVE has two loading matrices corresponding to the gene detection and gene count components, respectively, and therefore appears twice in Figure 5. In ZINB-WaVE, *π*_*ij*_ models whether a gene has been detected or not, and *μ*_*ij*_ models the mean for the read counts under negative binomial distribution. As in the previous section, we used the parameters 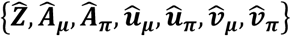 to replace the parameters 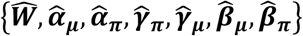 used in the original ZINB-WaVE paper for notational consistency:

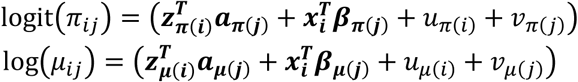

The loading matrix ***a***_***π***_ that models the gene detection component (π) is denoted as ZINB-WaVE_dropout_, and the loading matrix *a*_*μ*_ that models gene counts is denoted ZINB-WaVE_mean_.

### Trajectory inference

To evaluate the performance gains of scBFA in the context of trajectory inference, we used a recently published platform, dynverse, for which “gold standard” scRNA-seq datasets were already obtained and preprocessed, and scripts and performance metrics have already been defined to evaluate trajectory inference^32^. Gold standard datasets refer to those datasets in which experimental (non-computational) methods were used to annotate a dataset with trajectory information such as cell type clusters (“milestones”) and connections between cell type clusters (“milestone networks”). From the 27 datasets available on June 12, 2019 that met this gold standard status and were real (not synthetic), we filtered out datasets that had less than 170 cells, yielding a total of 20 benchmark datasets. As with our previous experiments, we used the HVG selection criterion to identify the top 2,000 varying genes for dimensionality reduction.

Our strategy for benchmarking trajectory inference was to identify an existing, top performing trajectory inference method that also uses dimensionality reduction in its pipeline, then replace that dimensionality reduction step with one of the methods we tested in our study. The dynverse paper identified Slingshot as a top performer^32^ To evaluate scBFA and the other methods, we substituted the PCA step of Slingshot with each dimensionality reduction method (scBFA, ZINB-WaVE, PCA, scImpute, SAVER, scrna2019, sctransform, scVI), and used dynverse to measure the performance of each modified version of Slingshot. The number of input latent dimensions was set to 10. Because the Slingshot implementation throws NA in cases where it is uncertain of the assignment of cells to a particular lineage, we removed two datasets from further evaluation because the number of NAs producted prevented calculation of the performance metrics (germline-human-both_guo.rds, mESC-differentiation_hayashi.rds). For each of 18 benchmarks (**Supplementary Table 8**), we used dynverse to compute three performance metrics with respect to the experimentally-gathered trajectory information: F1_milestones_, F1_branches_ and NMSE_lm_. F1_milestones_ measures the similarity between clustering membership of two trajectories. F1_branches_ compares the similarity between two branches’ assignment. NMSE_lm_ is a measurement of how well the position of a cell in the inferred trajectory predicts the position of the cell in the ground truth trajectory under linear regression. Larger values of F1_milestones_, F1_branches_ and NMSE_lm_ correspond to better performance. We obtained F1_milestones_, F1_branches_ and NMSE_lm_ via dynverse’s calculate_mapping and calculate_position_predict functions within the dyneval package, and converted raw values to ranks for Figure 6. The wrapper function to obtain Slingshots’ result is adapted from the internal function https://github.com/dynverse/ti_slingshot/blob/master/package/R/ti_slingshot.R.

### Visualization

After we obtain the embedding matrix from every method, we use the t-distributed stochastic embedding^44^ method to project the embedding matrix onto two dimensions for visualization as a scatterplot. In all visualizations, the number of factors used as input to t-SNE in each visualization is 10.

### Timing experiments

In the timing experiment of **Supplementary Figure 25**, we randomly subsampled 1k, 10k, 50k, 100k cells from the 1.3 Million 10x brain cell dataset from E18 mice and recorded the single-core execution time (in seconds) of all methods (PCA, ZINB-WaVE, scImpute, SAVER, sctransform, scrna2019 and Binary PCA) on the same machine. Due to the nonconvex nature of ZINB-WaVE’s objective function and different optimization scheme, we cannot strictly match the convergence criterion of ZINB-WaVE to scBFA. Therefore, we use the same number of iterations for each method that was used to generate the results in **Supplementary Figure 25**. Because scImpute requires specification of the number of cell clusters, we set the number of cell clusters to seven, similar to a previous study^45^ that used seven as an underestimate of true number of cell types.

