## Supplementary Materials for "scBFA: modeling detection patterns to mitigate technical noise in large-scale single cell genomics data"

Ruoxin Li[1,2], Gerald Quon[1,2,3,*]
[1]Graduate Group in Biostatistics, [2]Genome Center, [3]Department of Molecular and Cellular Biology, University of California, Davis, Davis, CA

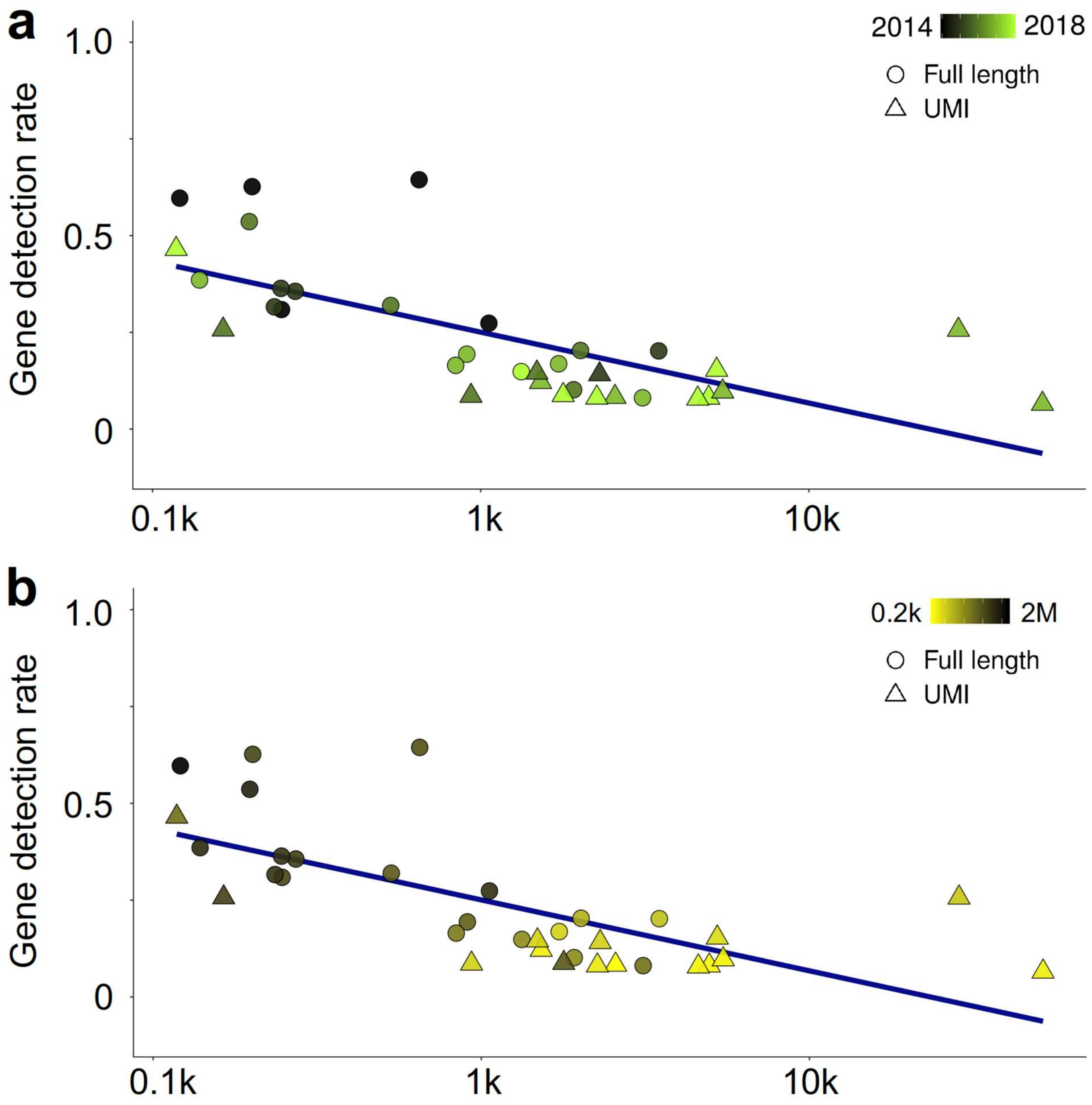

**Supplementary Figure 1: Gene detection rate decreases as the number of cells sequenced increases, under HVG selection.** (**a**) Gene detection rate as a function of the number of cells sequenced, across 36 scRNA-seq datasets (**Supplementary Table 1**). Datasets are colored by date of publication. (**b**) Same as (a), but studies are colored by library size.

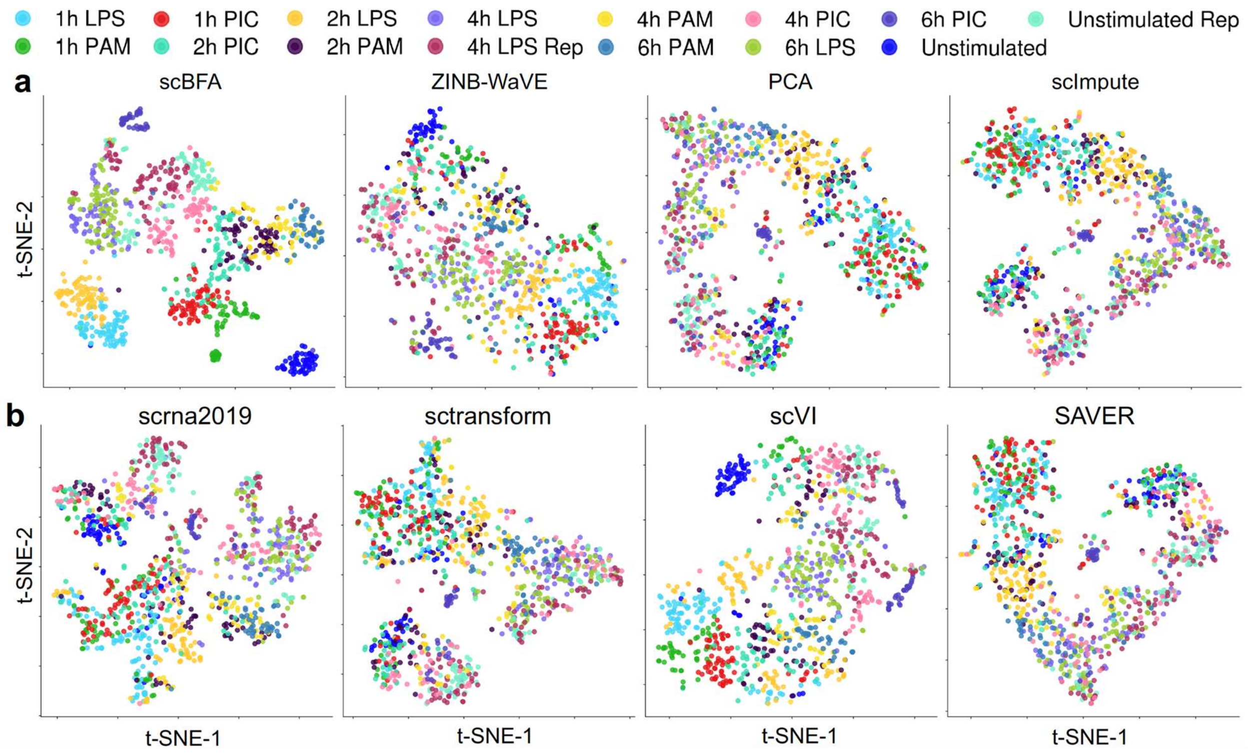

**Supplementary Figure 2: 2D tSNE visualization of the Dendritic benchmark of Shalek et al.** tSNE plots are generated based on the 10-dimensional embeddings learned by scBFA, PCA, ZINB-WaVE, scImpute, scrna2019, sctransform, scVI and SAVER. Cells are colored by their corresponding cell types.

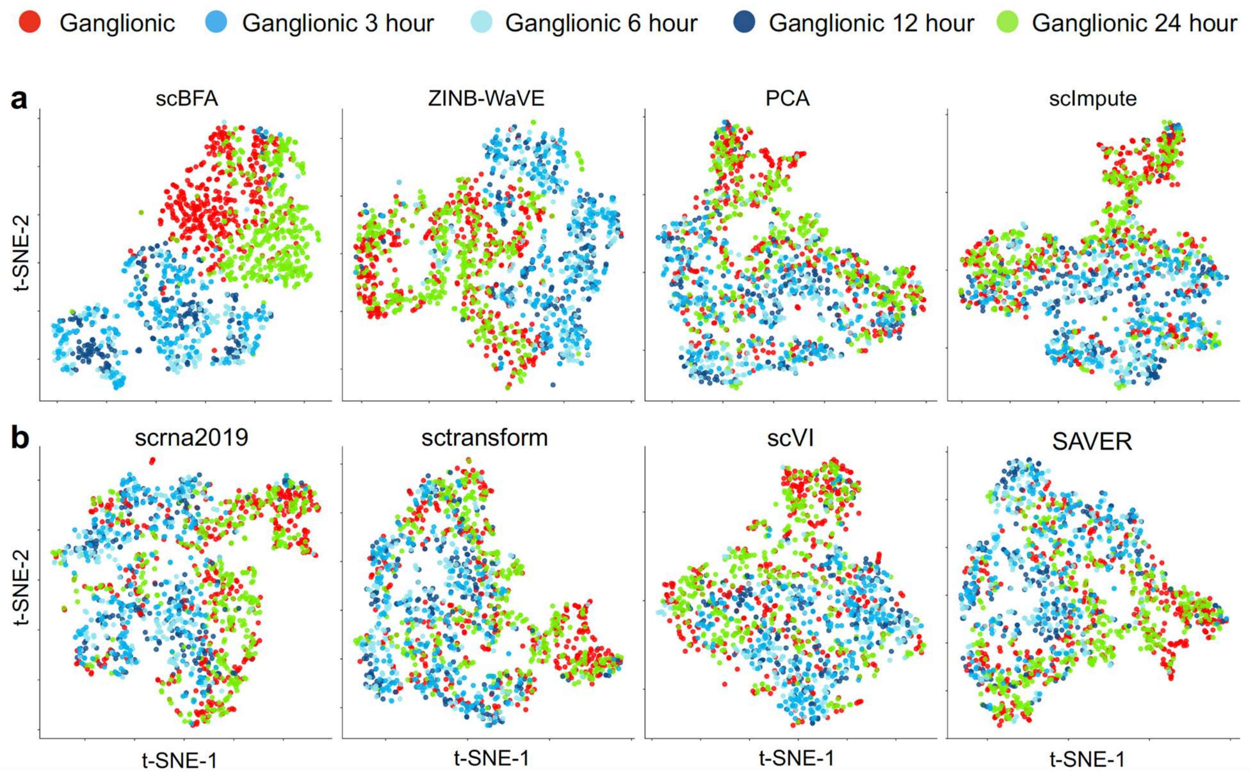

**Supplementary Figure 3: 2D tSNE visualization of the MGE benchmark of Mayer et al.** tSNE plots are generated based on the 10-dimensional embeddings learned by scBFA, PCA, ZINB-WaVE, scImpute, scrna2019, sctransform, scVI and SAVER. Cells are colored by their corresponding cell types.

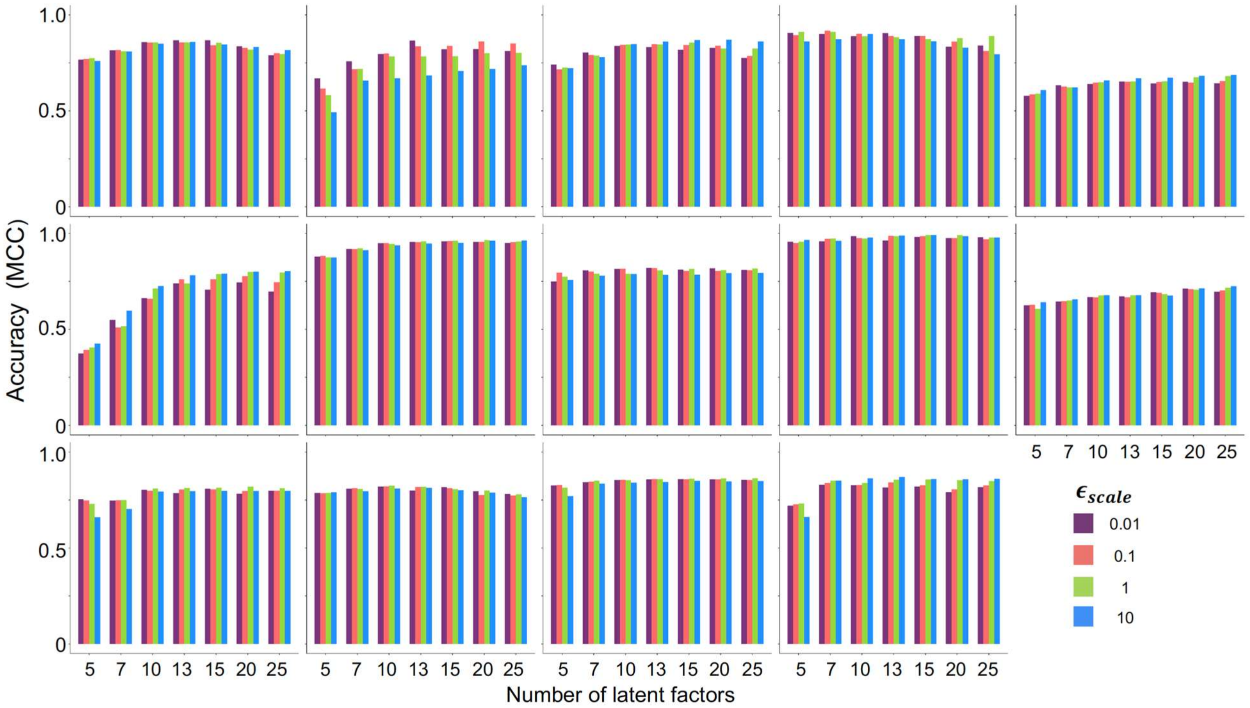

**Supplementary Figure 4: scBFA performance is robust with respect to selection of regularization parameters, under HVG selection.** Performance is measured via cross-validation of cell type classifiers trained on scRNA-seq benchmarks in the respective embedding spaces of scBFA under different regularization settings. Each color bar represents one selection of $\epsilon_{scale}$, indicating different levels of strength of regularization given to the learned model parameters. Here the universal regularization parameter $\epsilon = \epsilon_{scale} * \max(G, N)$, where we have set $\epsilon_{scale} \in \{0.01, 0.1, 1, 10\}$. Benchmarks from left to right, top to bottom: Dendritic, Pancreatic, DC, mESCs, HSPCs, MGE, Intestinal, MEM-T, H7-ESC, LSK, Myeloid, HSCs, PBMC, and LPS.

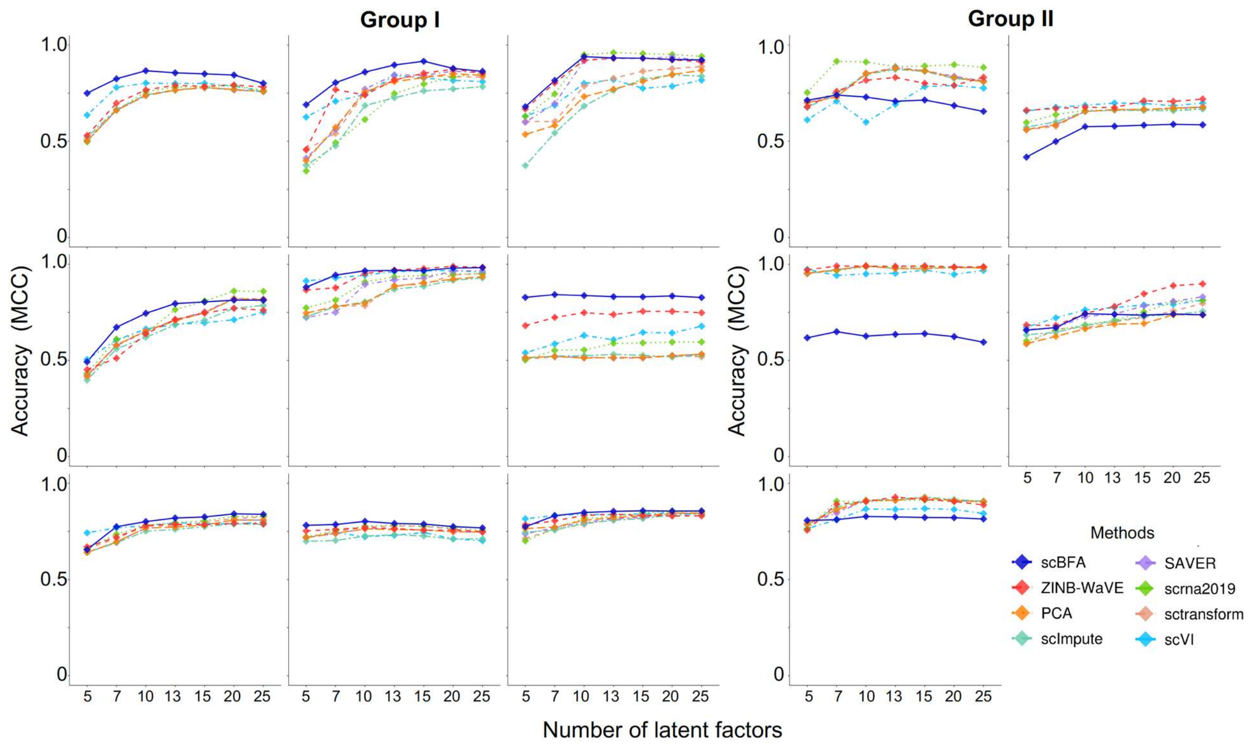

**Supplementary Figure 5: Performance of low dimensional embeddings with respect to cell type identification, under HEG gene selection.** Performance is measured via cross-validation of cell type classifiers trained on scRNA-seq benchmark data in the respective embedding spaces of each method, as a function of the number of latent dimensions specified. Benchmarks are grouped based on whether scBFA is a top performer (Group I) or performs poorly (Group II). The set of Group I benchmarks from left to right, top to bottom: Dendritic, Pancreatic, DC, MGE, Intestinal, MEM-T, Myeloid, HSCs and PBMC. The set of Group II benchmarks from left to right: mESCs, HSPC, H7-ESC, LSK and LPS.

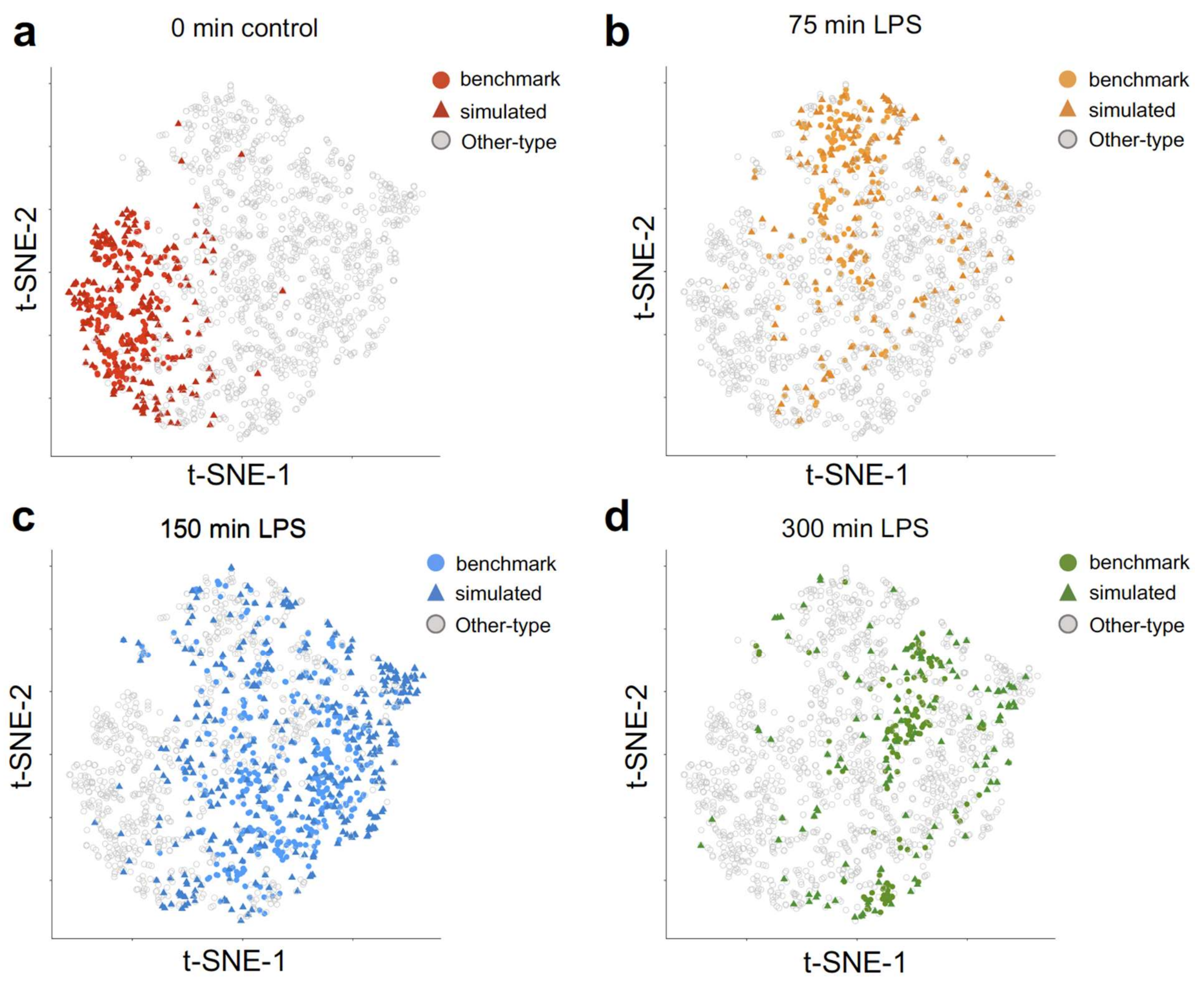

**Supplementary Figure 6: Simulation framework trained on the LPS benchmark generates simulated data that resemble the benchmark data.** The simulation framework was used to generate a simulated dataset of the same size as the LPS benchmark with HVG selection. The combined dataset (simulated and benchmark) were jointly projected into five dimensions via PCA, then drawn in 2D using tSNE. Cells are colored according to their corresponding cell type; the cell type of each simulated cell is based on the embedding parameter learned for the corresponding cell in the benchmark. Circles indicate cells from the benchmark, which triangles represent cells from the simulated dataset. Each plot illustrates the overlap of the same cell type across the benchmark and simulated datasets, while all other cell types are shown in grey. (**a**) 0 minute control. (**b**) 75 minutes post-LPS exposure. (**c**) 150 minutes post-LPS exposure. (**d**) 300 minutes post-LPS exposure.

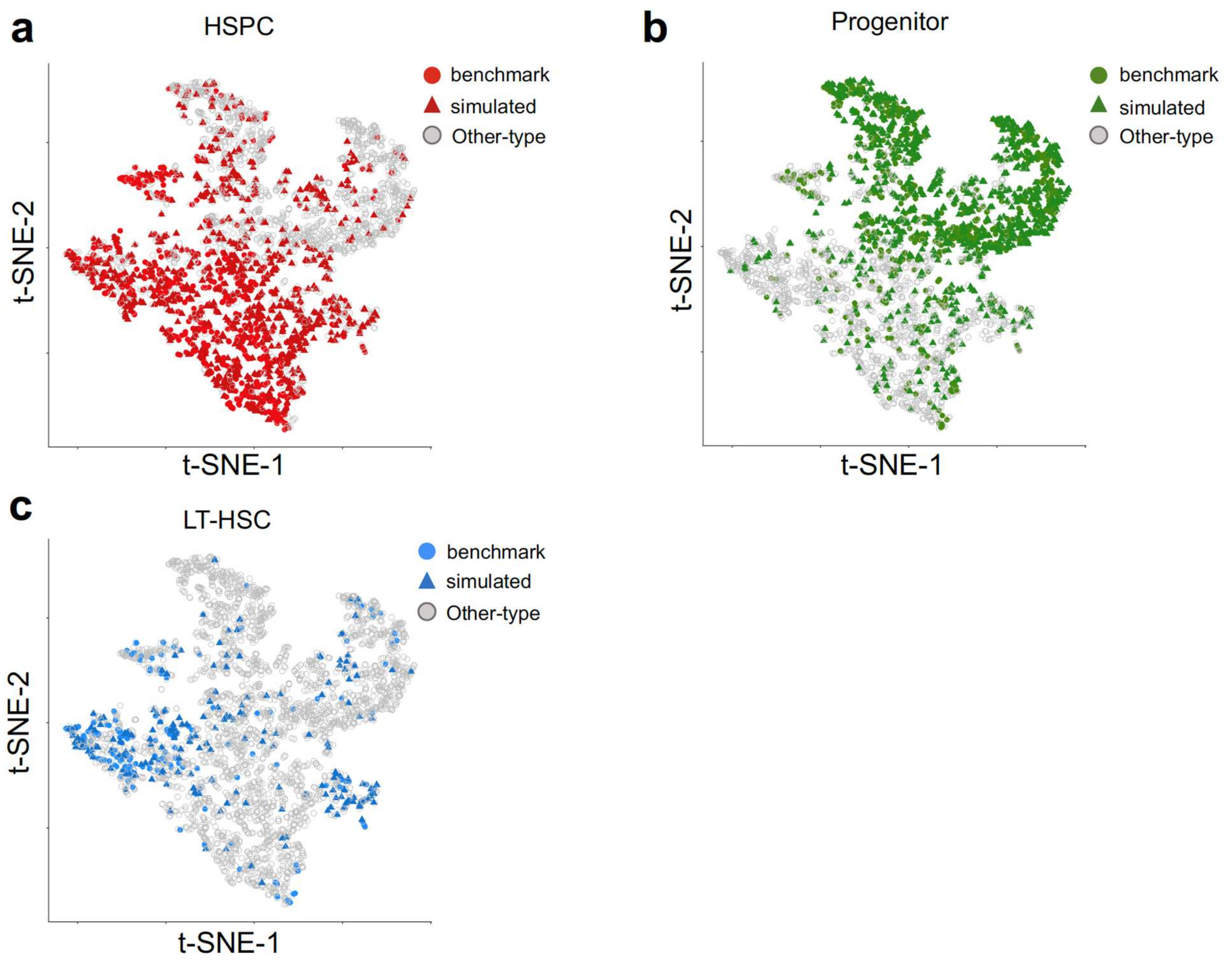

**Supplementary Figure 7: Simulation framework trained on the HSPC benchmark generates simulated data that resemble the benchmark data.** The simulation framework was used to generate a simulated dataset of the same size as the HSPC benchmark with HVG selection. The combined dataset (simulated and benchmark) were jointly projected into five dimensions via PCA, then drawn in 2D using tSNE. Cells are colored according to their corresponding cell type; the cell type of each simulated cell is based on the embedding parameter learned for the corresponding cell in the benchmark. Circles indicate cells from the benchmark, which triangles represent cells from the simulated dataset. Each plot illustrates the overlap of the same cell type across the benchmark and simulated datasets, while all other cell types are shown in grey. (**a**) HSPCs. (**b**) Progenitors. (**c**) Long term HSCs.

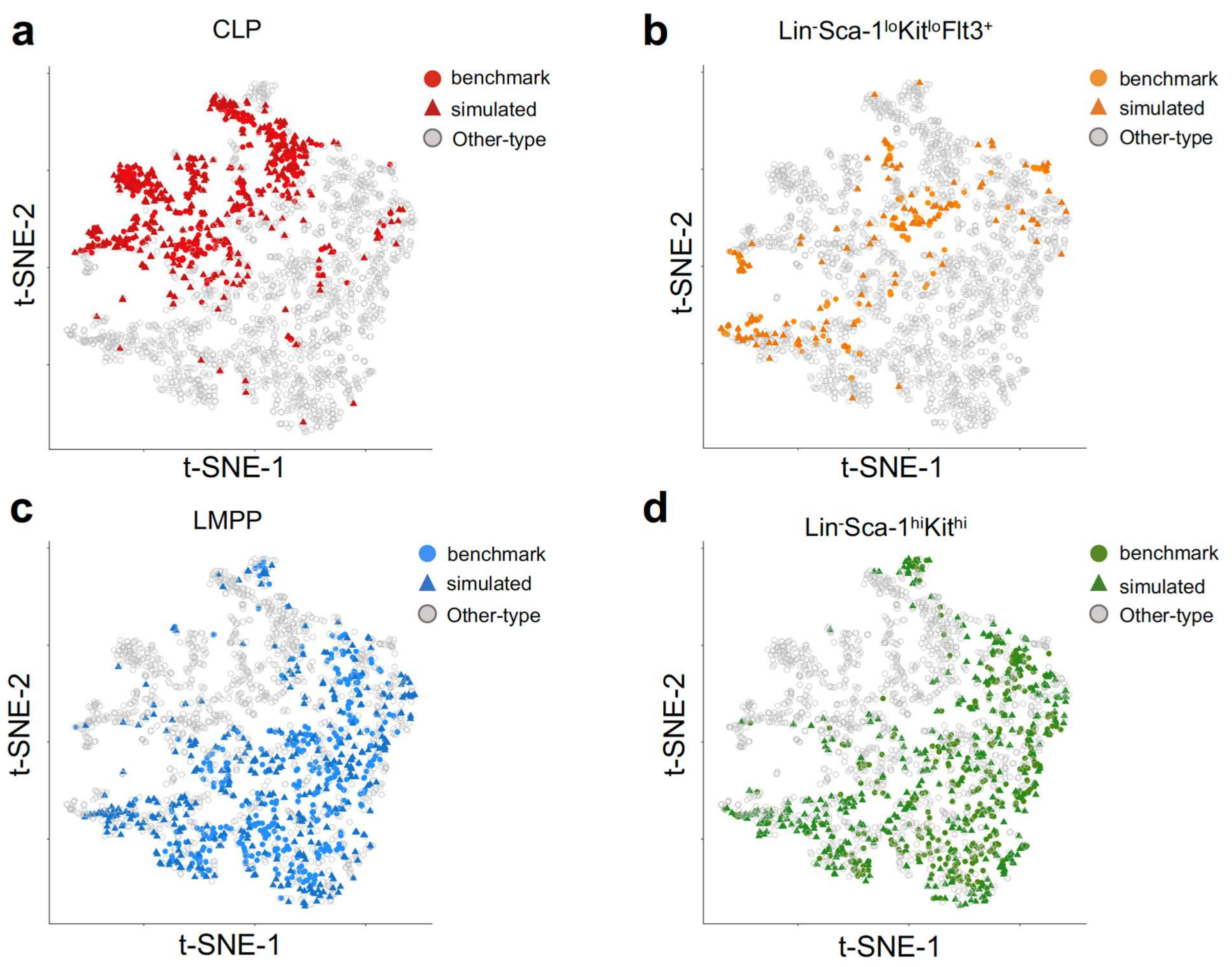

**Supplementary Figure 8: Simulation framework trained on the LSK benchmark generates simulated data that resemble the benchmark data.** The simulation framework was used to generate a simulated dataset of the same size as the LSK benchmark with HVG selection. The combined dataset (simulated and benchmark) were jointly projected into five dimensions via PCA, then drawn in 2D using tSNE. Cells are colored according to their corresponding cell type; the cell type of each simulated cell is based on the embedding parameter learned for the corresponding cell in the benchmark. Circles indicate cells from the benchmark, which triangles represent cells from the simulated dataset. Each plot illustrates the overlap of the same cell type across the benchmark and simulated datasets, while all other cell types are shown in grey. (**a**) Common lymphoid progenitors. (**b**) $Lin^{-}Sca\text{-}1^{lo}Kit^{lo}Flt3^{+}$ cells. (**c**) Lymphoid-primed multipotent progenitors. (**d**) $Lin^{-}Sca\text{-}1^{hi}Kit^{hi}$ cells.

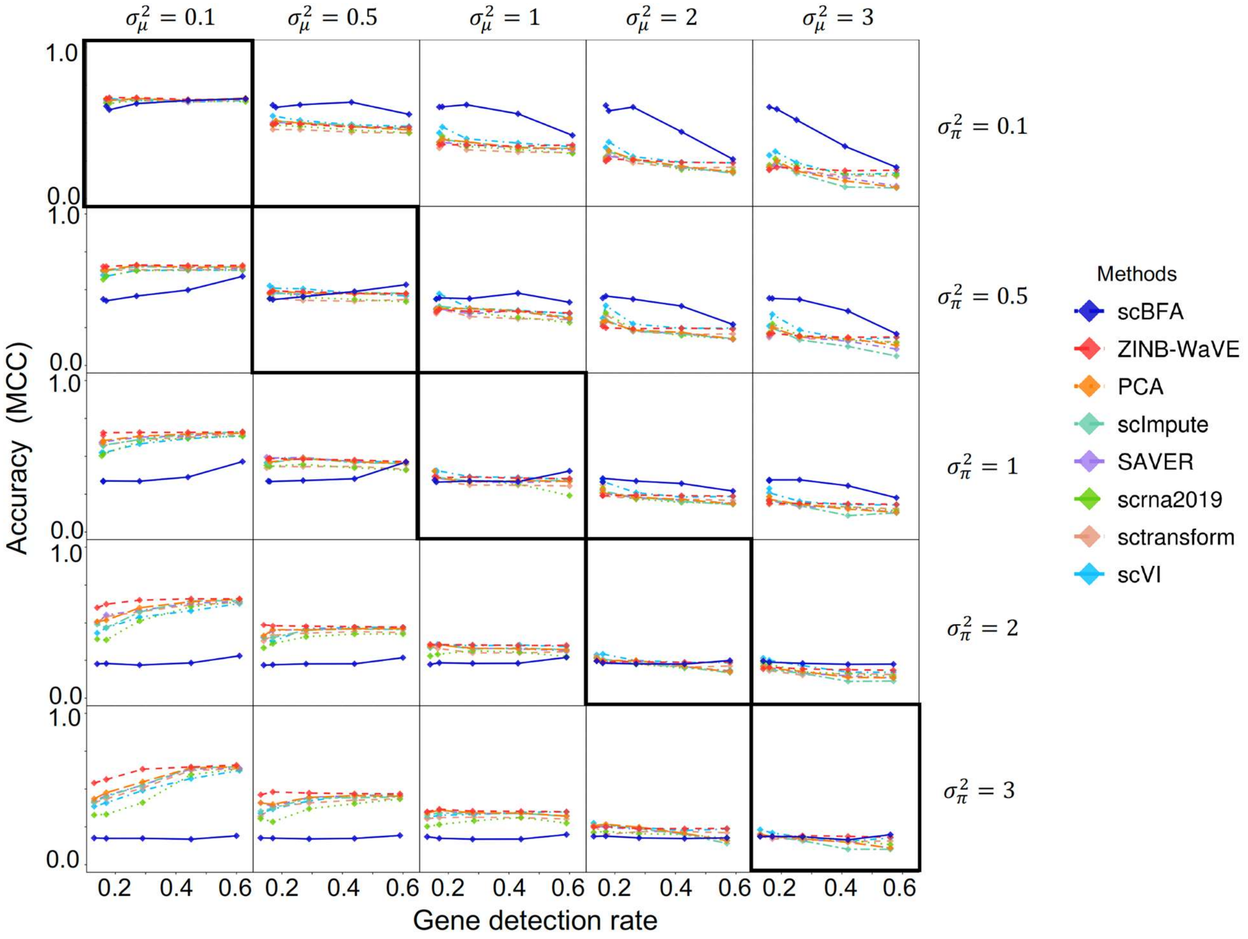

**Supplementary Figure 9: scBFA outperforms quantification models when the detection noise is smaller than the quantification noise (HVG, $r = 5$).** Rows represent different settings of (gene) detection noise ($\sigma_{\boldsymbol{\pi}}^2$), and columns represent different settings of (gene) quantification noise ($\sigma_\mu^2$). The diagonal represents simulations where the detection noise is equal to the quantification noise ($\sigma_\mu^2 = \sigma_\pi^2$), and the plots above the diagonal represent simulations where the detection noise is less than the quantification noise. The y-axis indicates the cross-validation performance of cell type predictors trained on embeddings learned from the simulated data, while the x-axis represents the gene detection rate that is manipulated by the parameter $\delta$. Here, the ground-truth embedding matrix is obtained by fitting ZINB-WaVE to the LPS benchmark under HVG selection. The dispersion parameter $r$ is set to be 5 in these simulations.

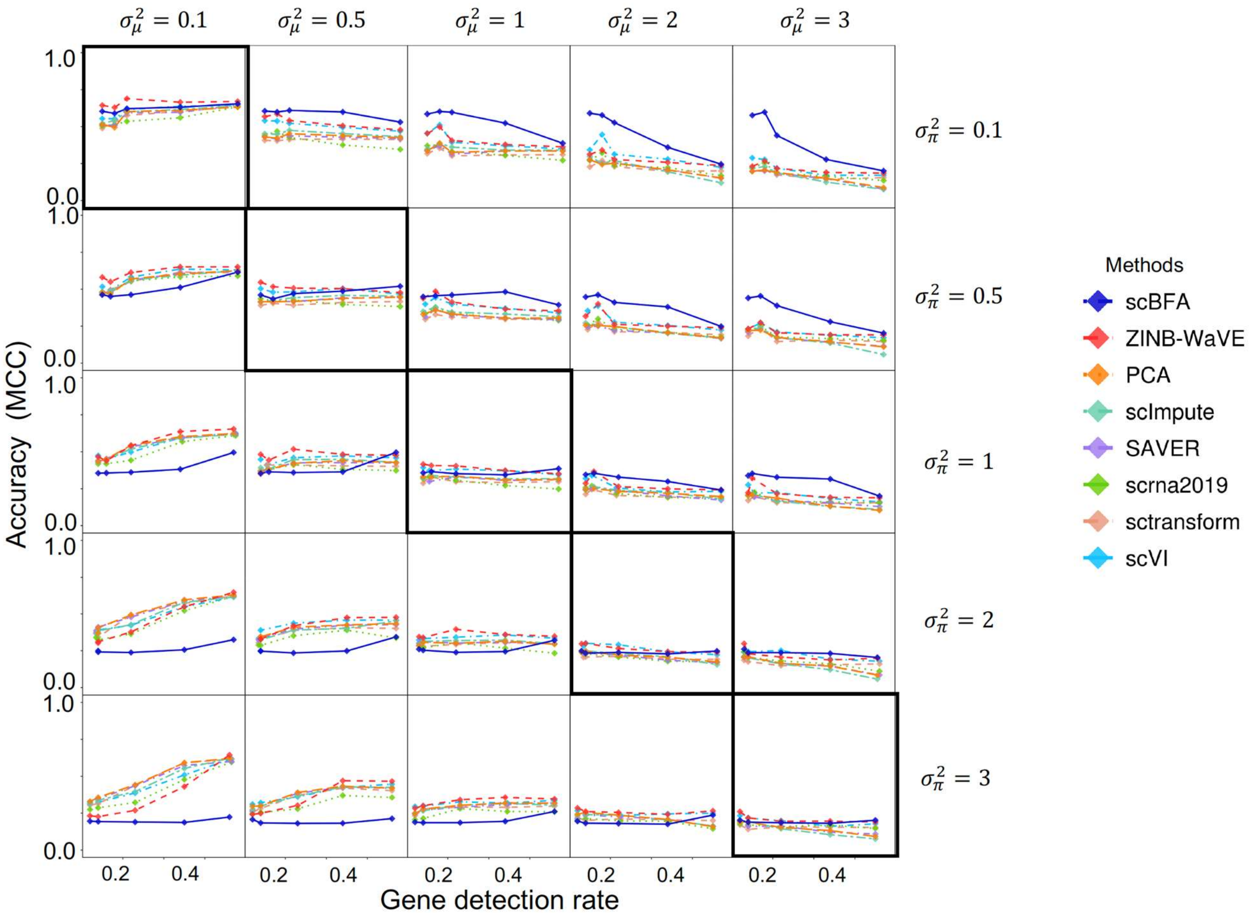

**Supplementary Figure 10: scBFA outperforms quantification models when the detection noise is smaller than the quantification noise (HVG, $r = 0.5$).** Rows represent different settings of (gene) detection noise ($\sigma_{\boldsymbol{\pi}}^2$), and columns represent different settings of (gene) quantification noise ($\sigma_\mu^2$). The diagonal represents simulations where the detection noise is equal to the quantification noise ($\sigma_\mu^2 = \sigma_\pi^2$), and the plots above the diagonal represent simulations where the detection noise is less than the quantification noise. The y-axis indicates the cross-validation performance of cell type predictors trained on embeddings learned from the simulated data, while the x-axis represents the gene detection rate that is manipulated by the parameter $\delta$. Here, the ground-truth embedding matrix is obtained by fitting ZINB-WaVE to the LPS benchmark under HVG selection. The dispersion parameter $r$ is set to be 0.5 in these simulations.

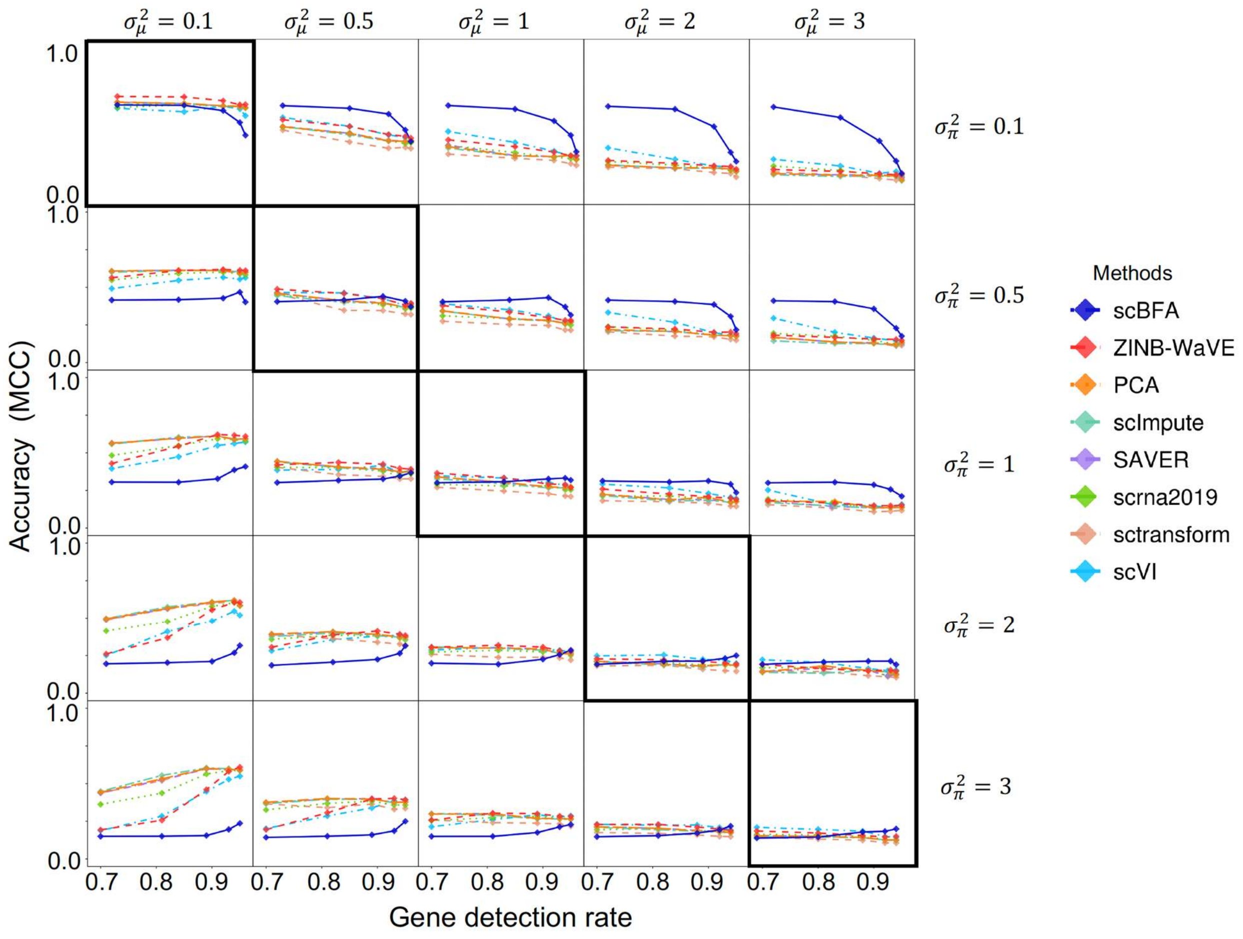

**Supplementary Figure 11: scBFA outperforms quantification models when gene detection noise is smaller than quantification noise (HEG, $r = 1$).** Rows represent different settings of (gene) detection noise ($\sigma_{\boldsymbol{\pi}}^2$), and columns represent different settings of (gene) quantification noise ($\sigma_\mu^2$). The diagonal represents simulations where the detection noise is equal to the quantification noise ($\sigma_\mu^2 = \sigma_\pi^2$), and the plots above the diagonal represent simulations where the detection noise is less than the quantification noise. The y-axis indicates the cross-validation performance of cell type predictors trained on embeddings learned from the simulated data, while the x-axis represents the gene detection rate that is manipulated by the parameter $\delta$. Here, the ground-truth embedding matrix is obtained by fitting ZINB-WaVE to the LPS benchmark under HEG selection. The dispersion parameter $r$ is set to be 1 in these simulations.

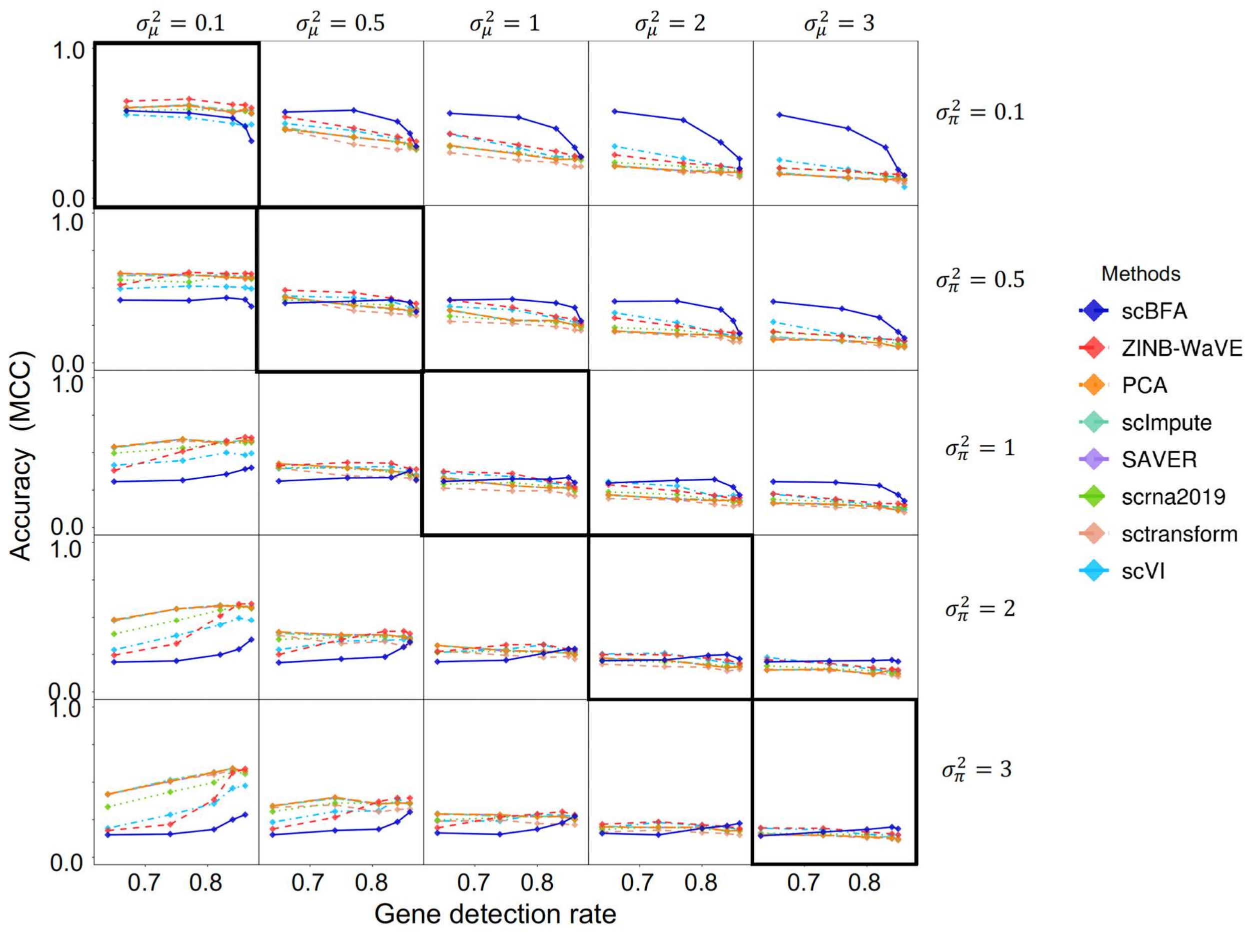

**Supplementary Figure 12: scBFA outperforms quantification models when gene detection noise is smaller than quantification noise (HEG, $r = 0.5$).** Rows represent different settings of (gene) detection noise ($\sigma_{\pi}^2$), and columns represent different settings of (gene) quantification noise ($\sigma_\mu^2$). The diagonal represents simulations where the detection noise is equal to the quantification noise ($\sigma_\mu^2 = \sigma_\pi^2$), and the plots above the diagonal represent simulations where the detection noise is less than the quantification noise. The y-axis indicates the cross-validation performance of cell type predictors trained on embeddings learned from the simulated data, while the x-axis represents the gene detection rate that is manipulated by the parameter $\delta$. Here, the ground-truth embedding matrix is obtained by fitting ZINB-WaVE to the LPS benchmark under HEG selection. The dispersion parameter $r$ is set to be 0.5 in these simulations.

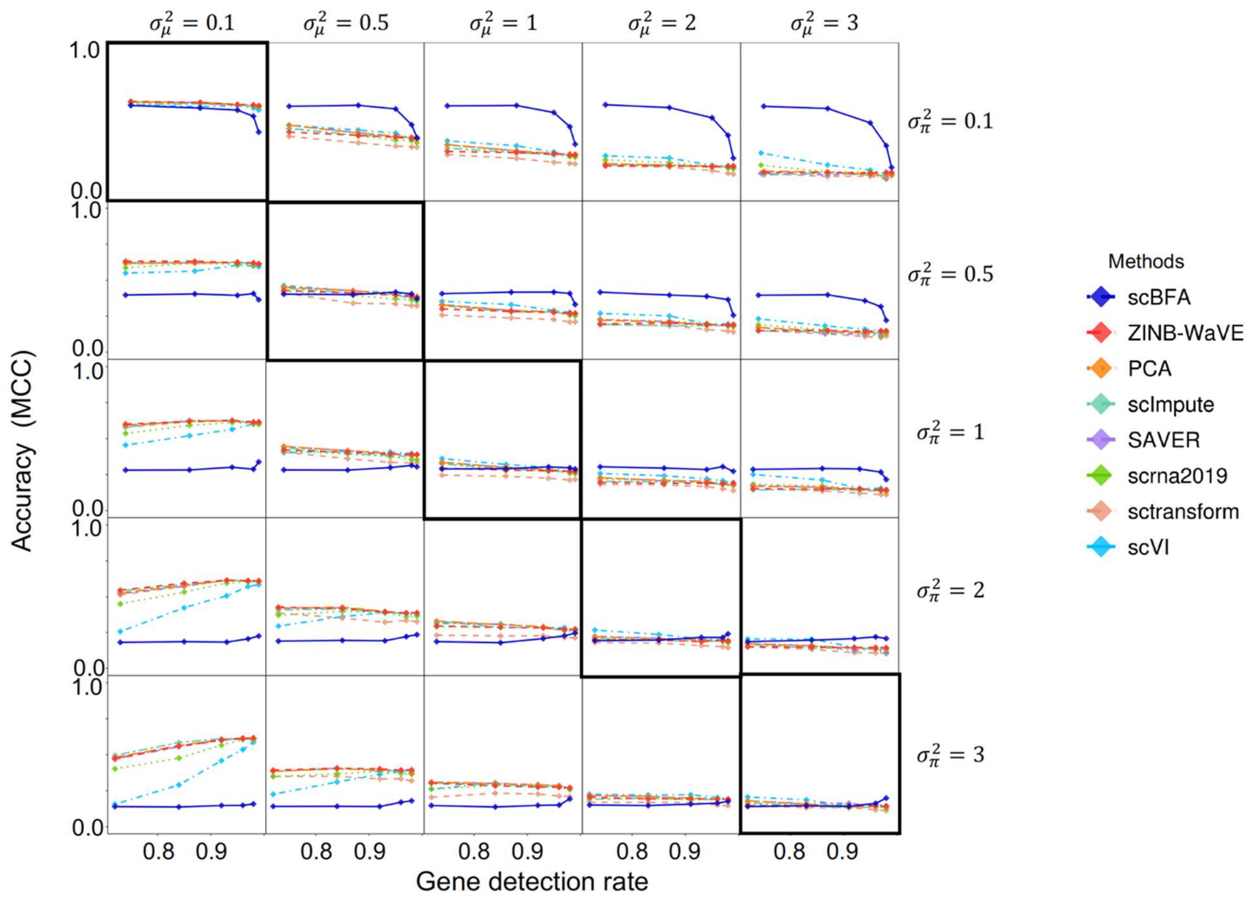

**Supplementary Figure 13: scBFA outperforms quantification models when gene detection noise is smaller than quantification noise (HEG, $r = 5$).** Rows represent different settings of (gene) detection noise ($\sigma_{\boldsymbol{\pi}}^2$), and columns represent different settings of (gene) quantification noise ($\sigma_\mu^2$). The diagonal represents simulations where the detection noise is equal to the quantification noise ($\sigma_\mu^2 = \sigma_\pi^2$), and the plots above the diagonal represent simulations where the detection noise is less than the quantification noise. The y-axis indicates the cross-validation performance of cell type predictors trained on embeddings learned from the simulated data, while the x-axis represents the gene detection rate that is manipulated by the parameter $\delta$. Here, the ground-truth embedding matrix is obtained by fitting ZINB-WaVE to the LPS benchmark under HEG selection. The dispersion parameter $r$ is set to be 5 in these simulations.

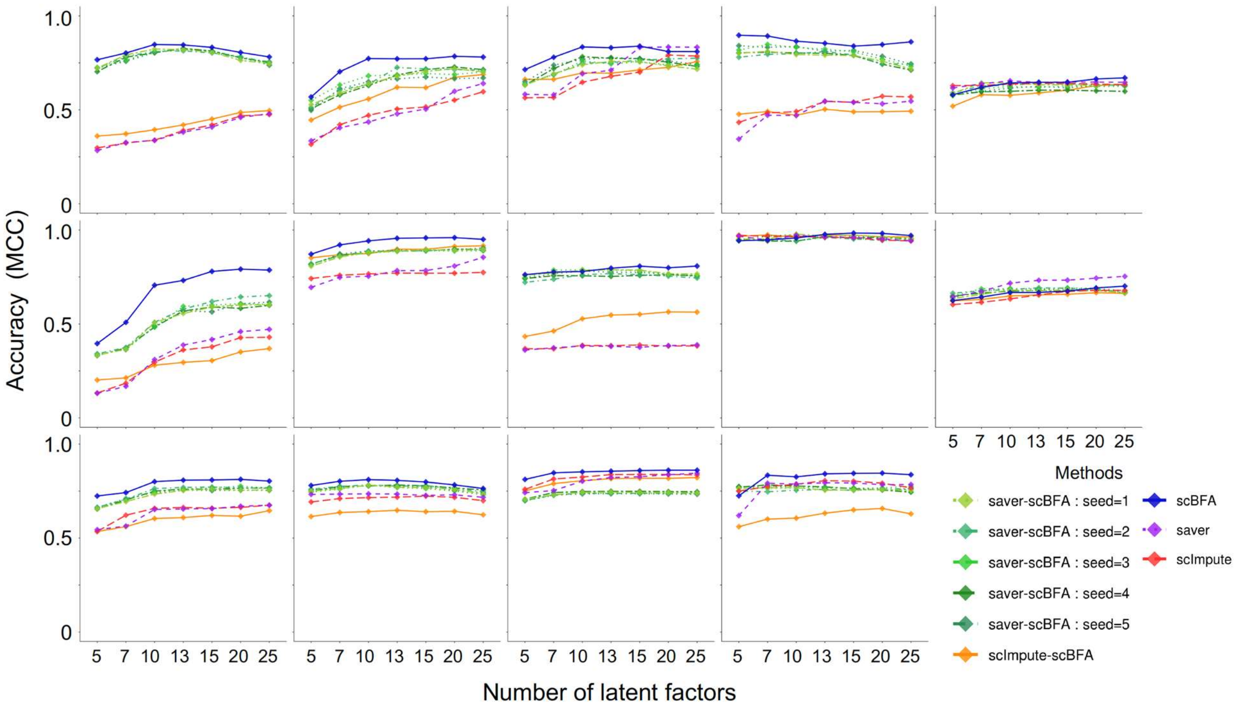

**Supplementary Figure 14: scBFA trained on the observed count matrix outperforms scBFA trained on an imputed count matrix, with respect to cell type identification.** Performance is measured via cross-validation of cell type classifiers trained on scRNA-seq benchmark data in the respective embedding spaces of each method, as a function of the number of latent dimensions specified. Across all 14 benchmarks, scBFA performs best without imputation compared to scBFA trained on an imputed expression matrix generated from either SAVER or scImpute.

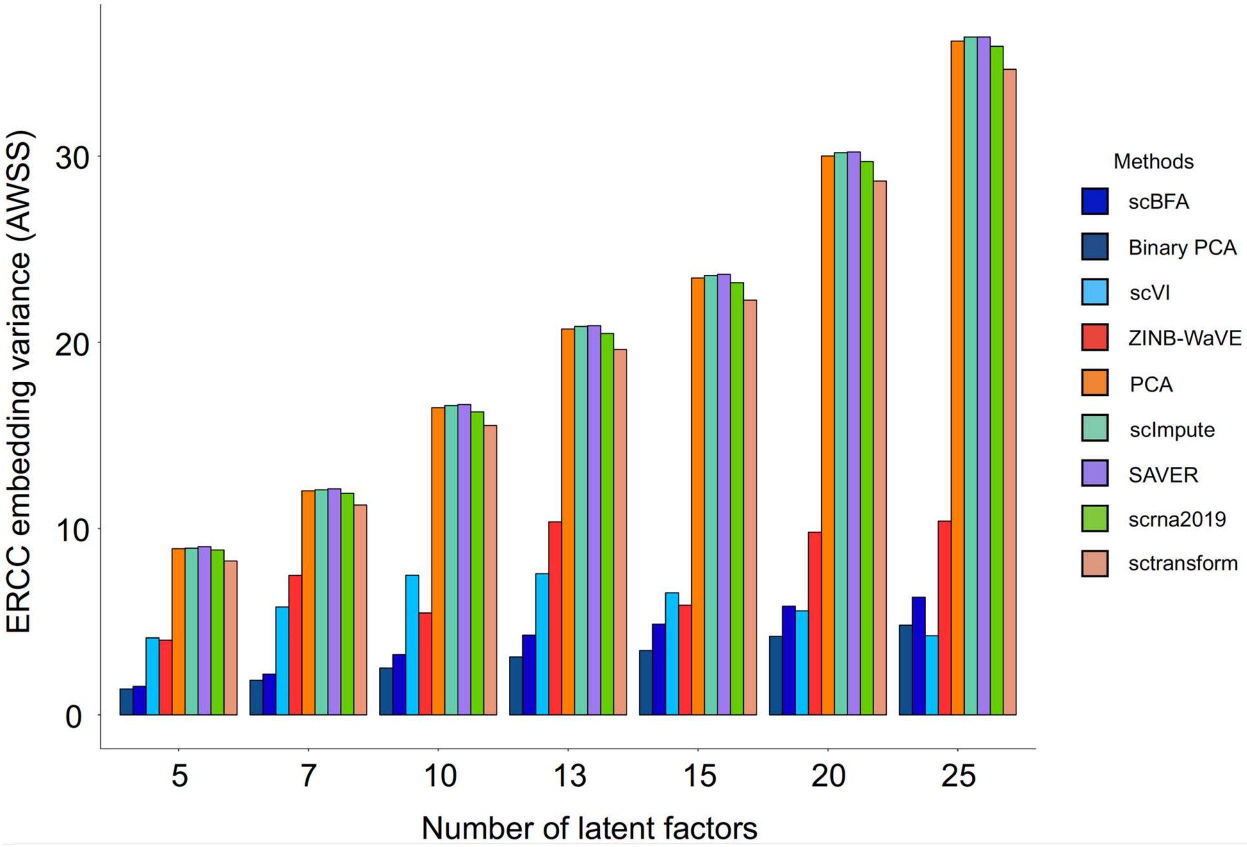

**Supplementary Figure 15: Comparison of the variance in embedding dimensions of the ERCC dataset of Zheng et al.** Methods are compared based on the variance of the embeddings learned over the ERCC dataset generated by Zheng et al. Variance is measured as AWSS (Average Within-Group Sum of Squares), as a function of the number of latent dimensions specified. Gene detection models (scBFA and Binary PCA) yield embeddings with lowest variance.

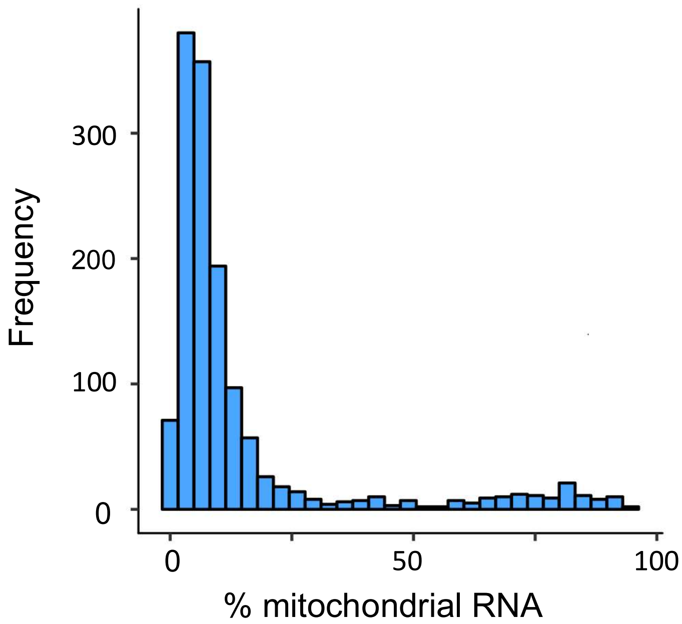

**Supplementary Figure 16: Distribution of the fraction of reads of each cell mapping to mitochondrial genes in the Dendritic benchmark of Shalek et al.** The fraction is calculated based on the HEG selection criterion.

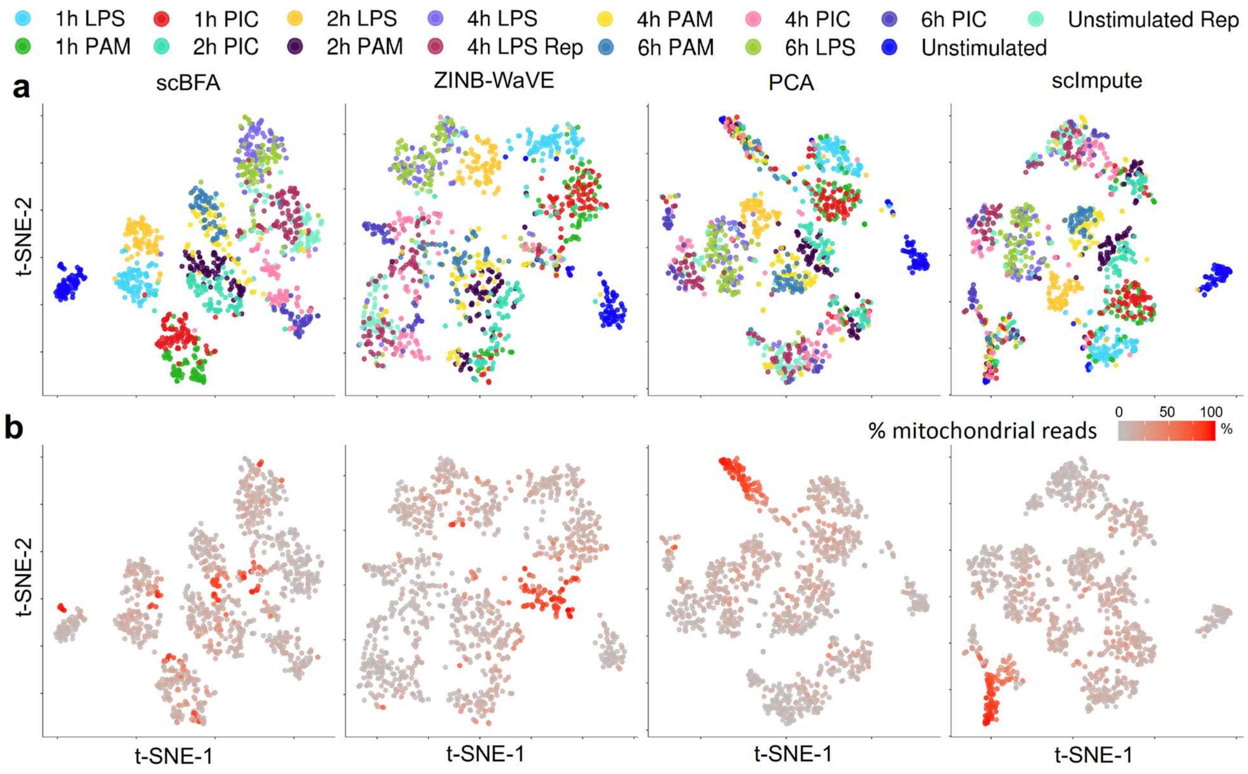

**Supplementary Figure 17: 2D tSNE visualization of the Dendritic benchmark of Shalek et al.** (**a**) 2D tSNE visualization of the 10-dimensional embedding generated by scBFA, ZINB-WaVE, PCA and scImpute on the Dendritic benchmark under HEG selection, when cells with high mitochondrial RNA content are kept in the analysis. Cells are colored according to their corresponding cell types and states**.** (**b**) Same visualization as in (a), but cells are colored according to the fraction of their reads that map to mitochondrial genes.

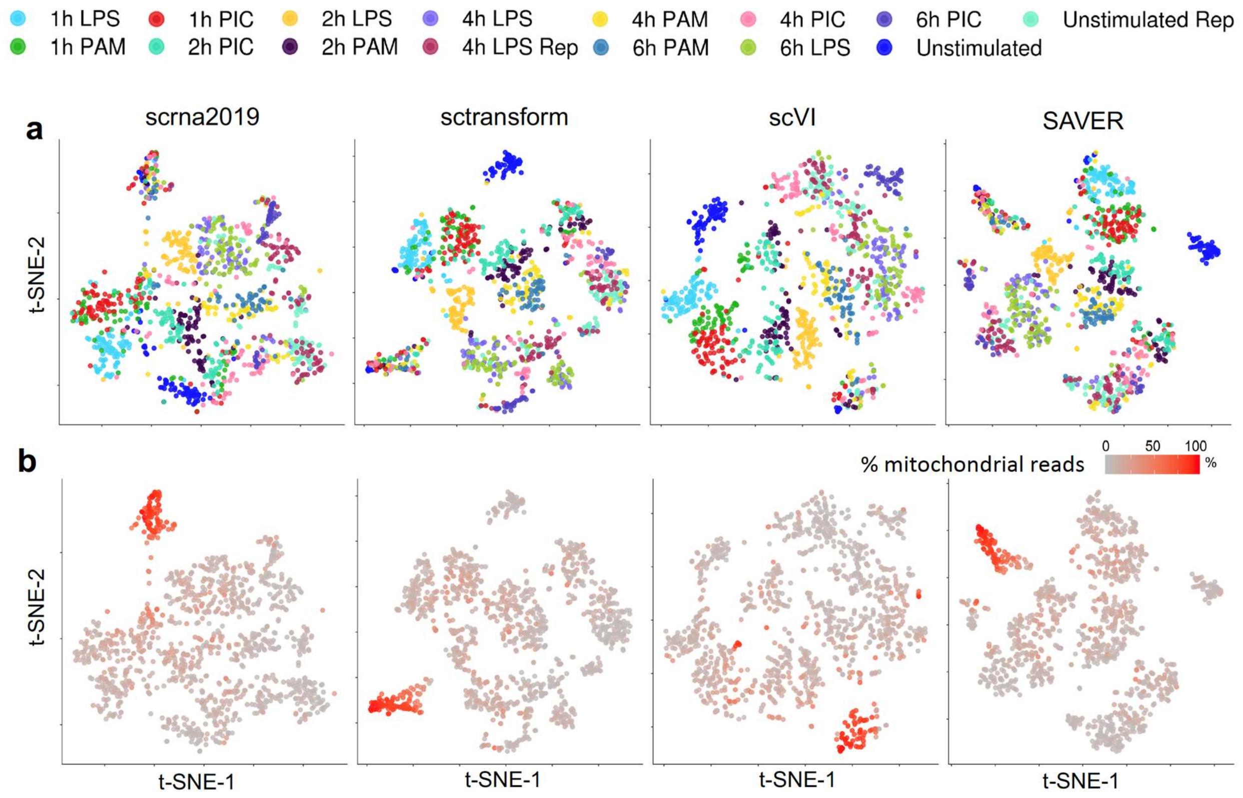

**Supplementary Figure 18: 2D tSNE visualization of the Dendritic benchmark of Shalek et al.** (**a**) 2D tSNE visualization of the 10-dimensional embedding generated by scrna2019, sctransform, scVI and SAVER on the Dendritic benchmark under HEG selection, when cells with high mitochondrial RNA content are kept in the analysis. Cells are colored according to their corresponding cell types and states**.** (**b**) Same visualization as in (a), but cells are colored according to the fraction of their reads that map to mitochondrial genes.

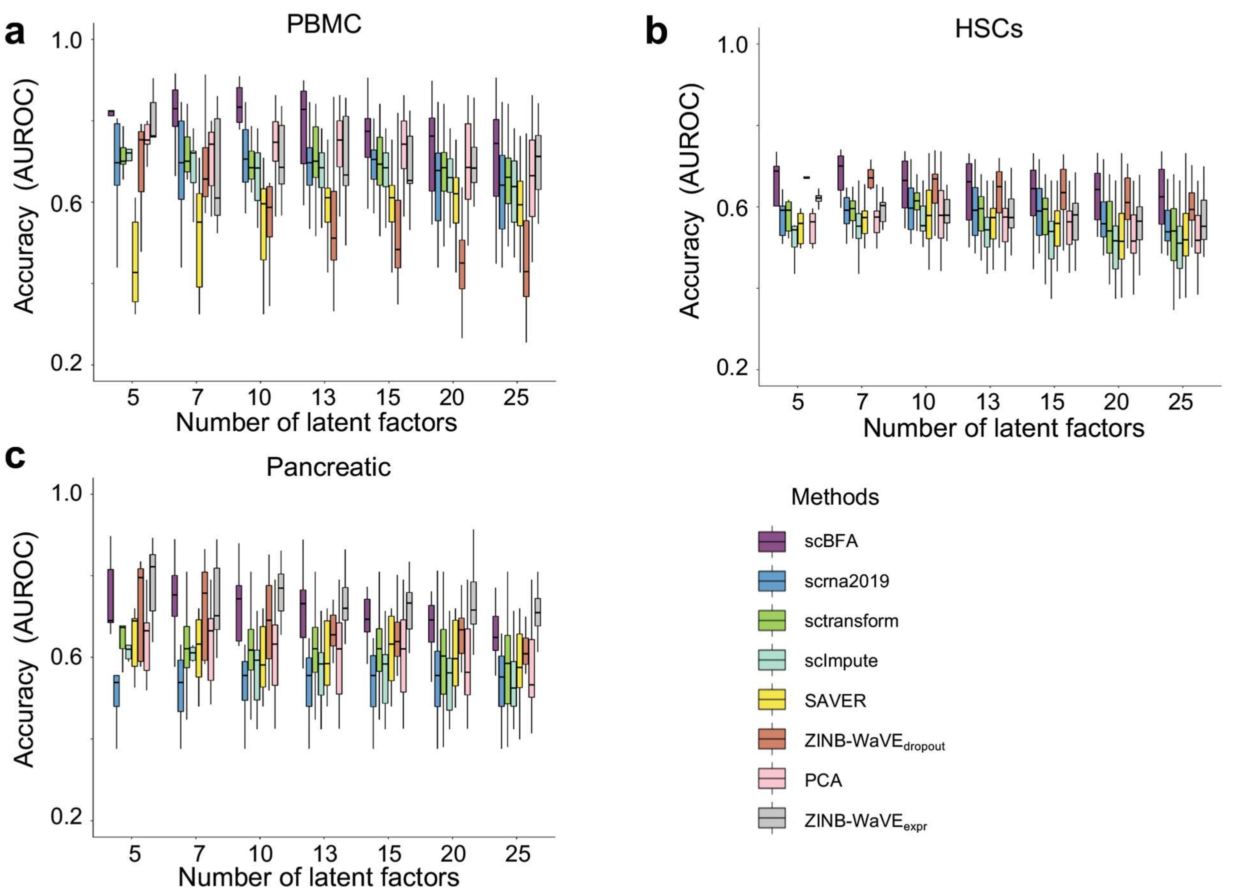

**Supplementary Figure 19: scBFA is better informed by cell type markers than quantification models, under HEG selection.** Each latent factor learned from each method was evaluated based on how much influence established cell type markers had on its embeddings, as measured by the area under the curve (AUROC) metric. Each boxplot represents the AUROC of all latent factors for a given method, for a given benchmark. ZINB-WaVE is represented twice, once for the latent dimensions inferred by their gene detection pattern (ZINB-WaVE$_{dropout}$), and once for the latent dimensions inferred from the gene counts (ZINB-WaVE$_{expr}$). (**a**) PBMC benchmark. (**b**) HSCs benchmark. (**c**) Pancreatic benchmark.

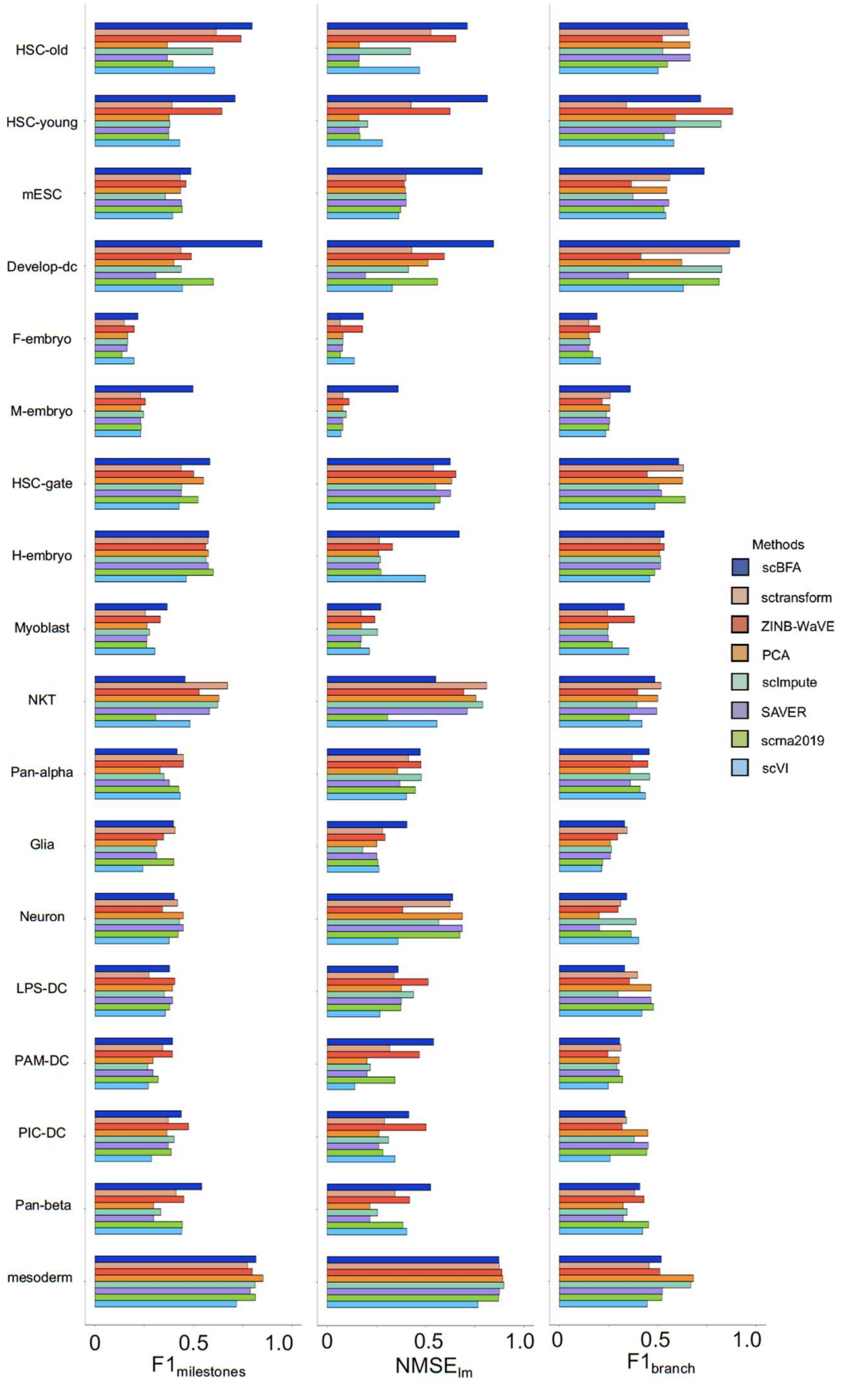

**Supplementary Figure 20: scBFA improves trajectory inference of the method Slingshot compared to other dimensionality reduction methods.** The y-axis represents the set of 18 "gold standard" trajectory inference benchmarks from Dynverse. These benchmarks are processed under HVG gene selection. For each dimensionality reduction method, the trajectory inference method Slingshot was adapted by replacing its PCA step with a corresponding dimensionality reduction method, and the resulting performance evaluated. Performance here is measured via the $F1_{milestone}$, $NMSE_{lm}$ and $F1_{branch}$ scores that measure how well the inferred trajectory matches the ground truth provided by Dynverse. $F1_{milestone}$ and $F1_{branch}$ are based on the quality of clustering of cells in the trajectory, while $NMSE_{lm}$ assesses how well the position of a cell in the inferred trajectory predicts the position of the cell in the ground truth trajectory.

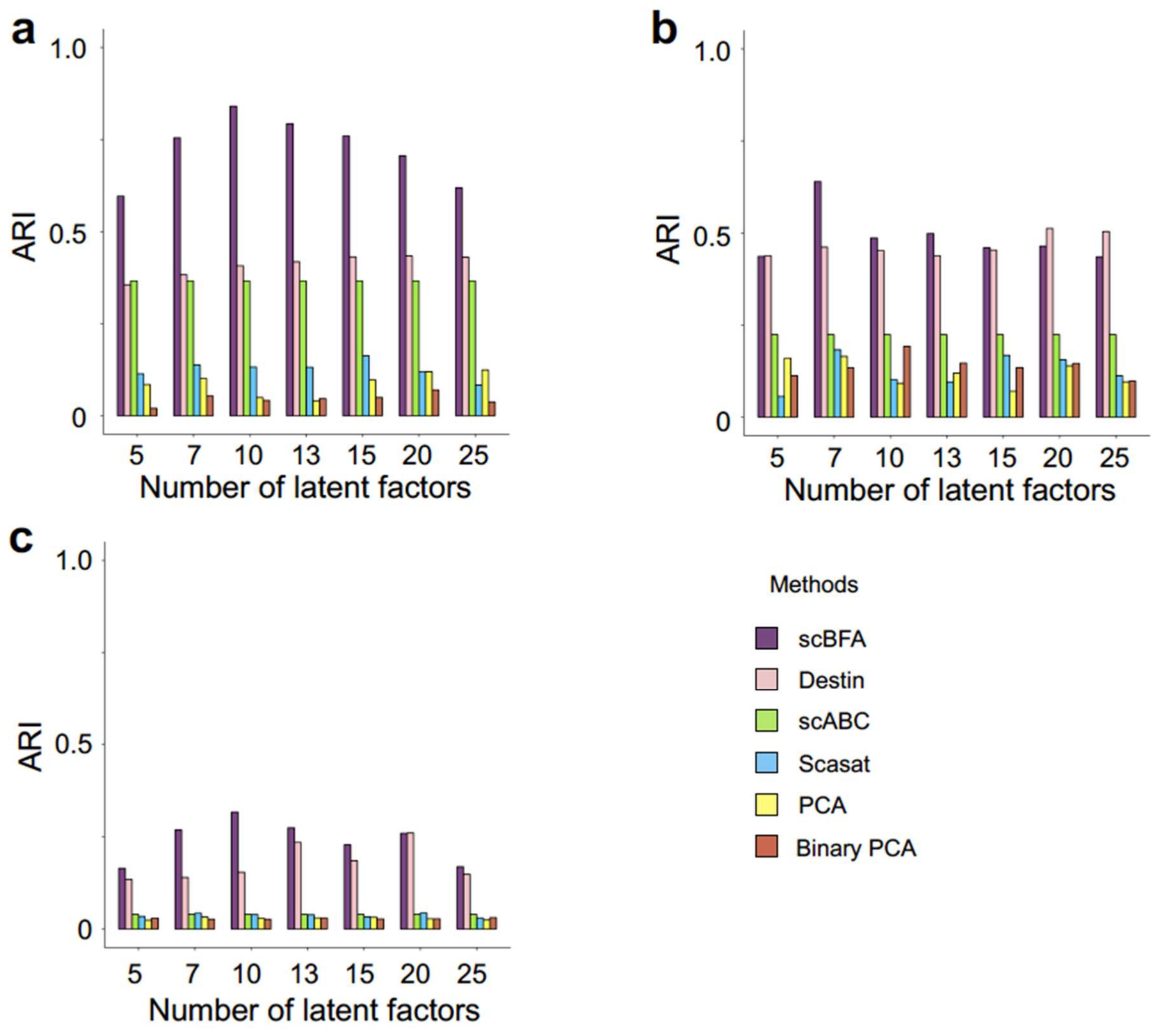

**Supplementary Figure 21: scBFA accurately recovers cell type identity in scATAC-seq benchmarks.** ARI (Adjusted Rand Index) measures the clustering accuracy of different methods on each scATAC-seq benchmark, as a function of the number of latent dimensions specified. The benchmarks from left to right, top to bottom are GSE96769, GSE74310 and GSE107816.

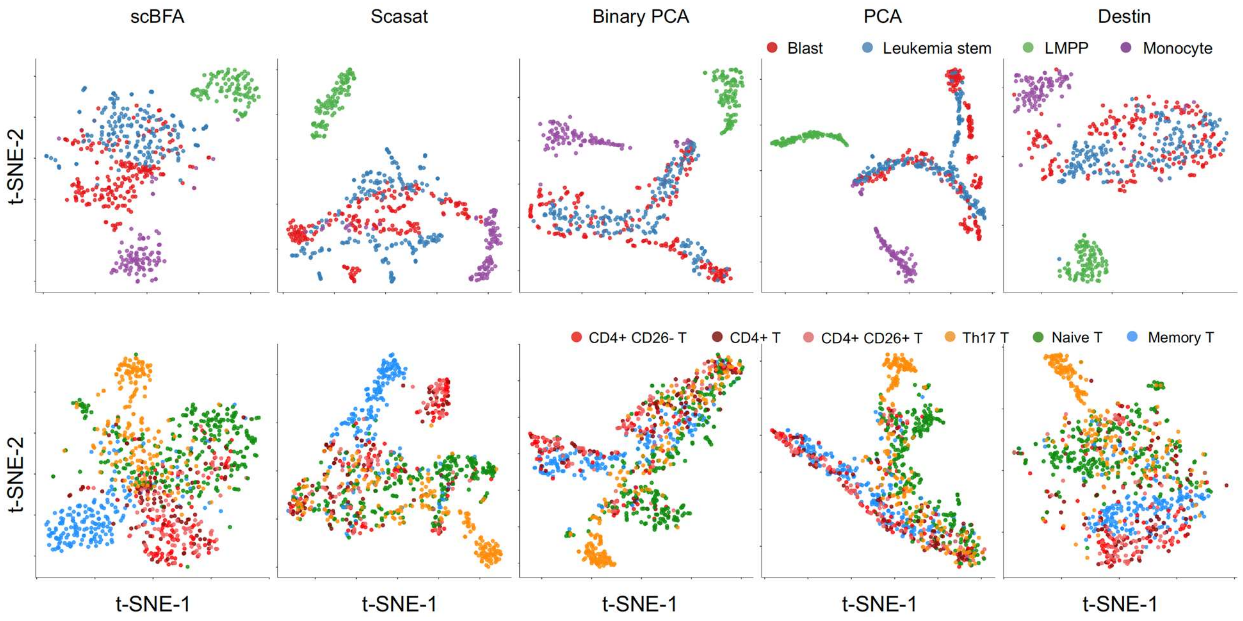

**Supplementary Figure 22: 2D tSNE visualization of the scATAC-seq datasets GSE74310 and GSE107816.** tSNE plots are generated based on the 10-dimensional embeddings learned by scBFA, Scasat, Binary PCA, PCA and Destin. The upper panel represents GSE74310 and the lower panel represents GSE107816**.** Cells are colored by their corresponding cell types.

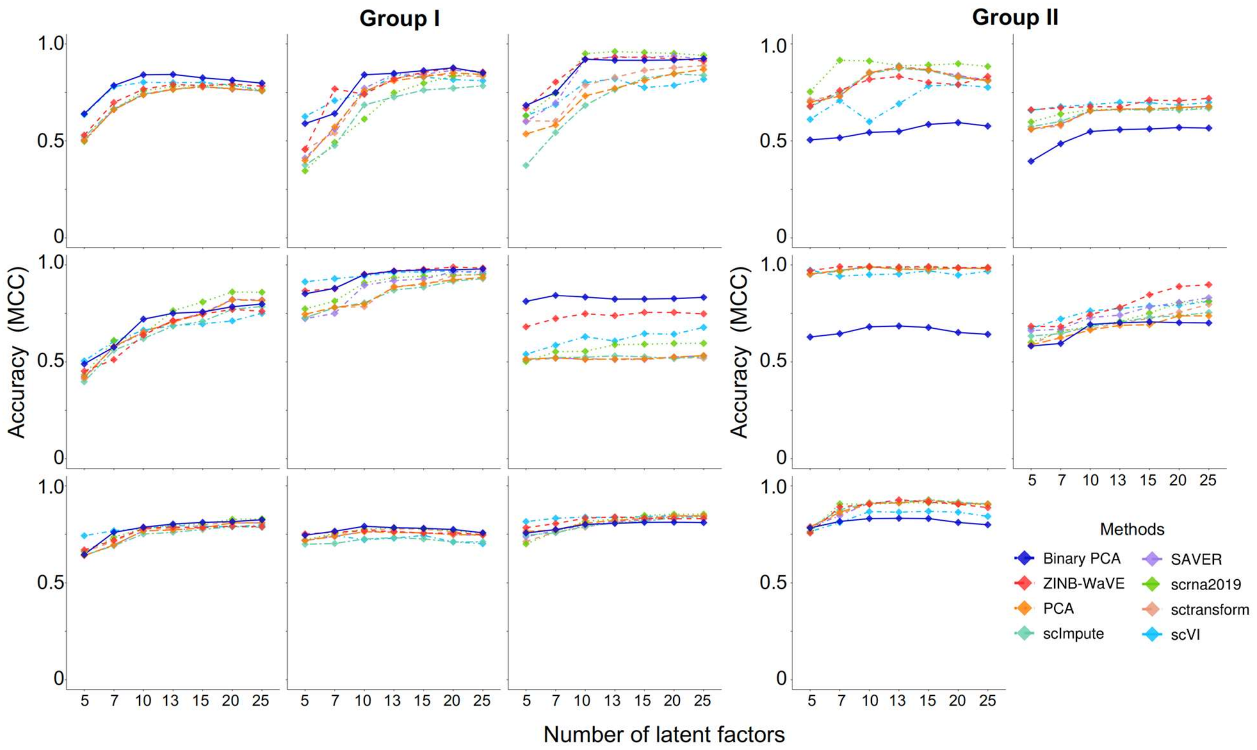

**Supplementary Figure 23: Binary PCA performance versus quantification models under HEG selection.** Performance is measured via cross-validation of cell type classifiers trained on scRNA-seq benchmark data in the respective embedding spaces of each method, as a function of the number of latent dimensions specified. Benchmarks are grouped based on whether scBFA is a top performer (Group I) or performs poorly (Group II). The set of Group I benchmarks from left to right, top to bottom: Dendritic, Pancreatic, DC, MGE, Intestinal, MEM-T, Myeloid, HSCs, PBMC. The set of Group II benchmarks from left to right: mESCs, HSPC, H7-ESC, LSK and LPS.

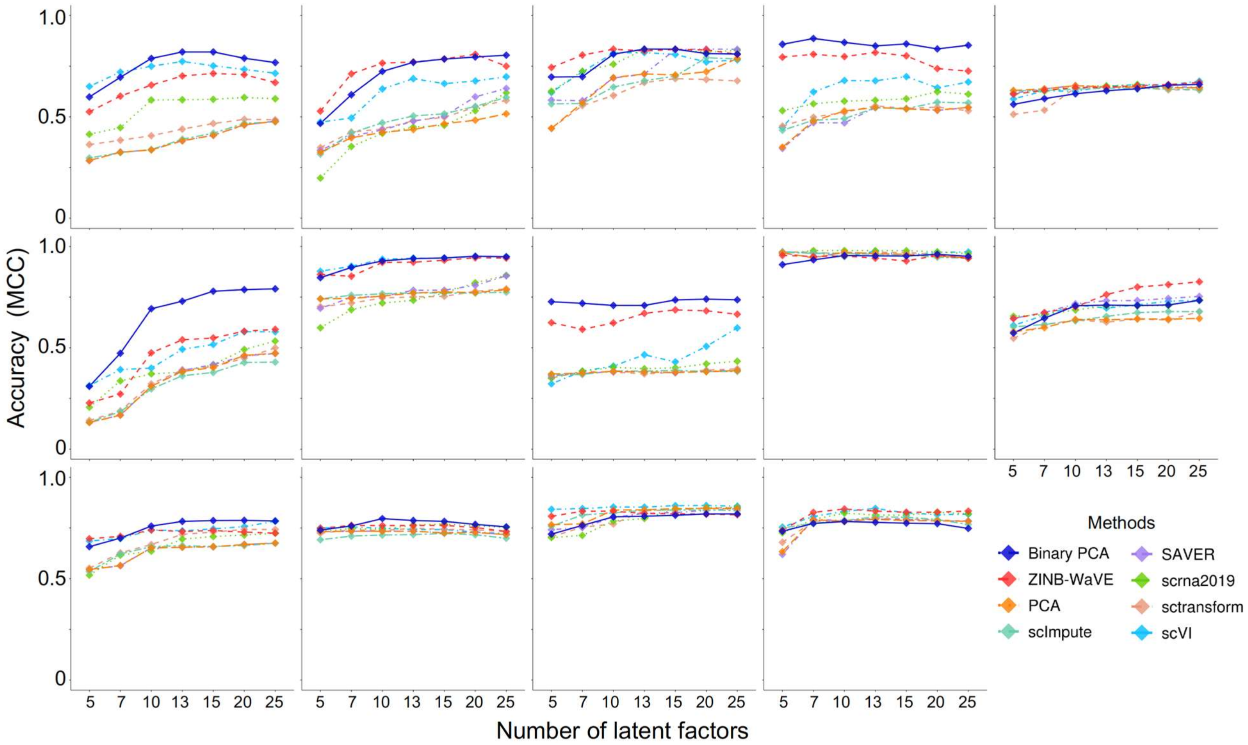

**Supplementary Figure 24: Binary PCA performance versus quantification models under HVG selection.** Cross-validation performance of cell type classifiers trained on low dimensional embeddings of scRNA-seq benchmark data, as a function of the number of latent dimensions specified. Benchmarks from left to right, top to bottom: Dendritic, Pancreatic, DC, mESCs, HSPCs, MGE, Intestinal, MEM-T, H7-ESC, LSK, Myeloid, HSCs, PBMC, and LPS.

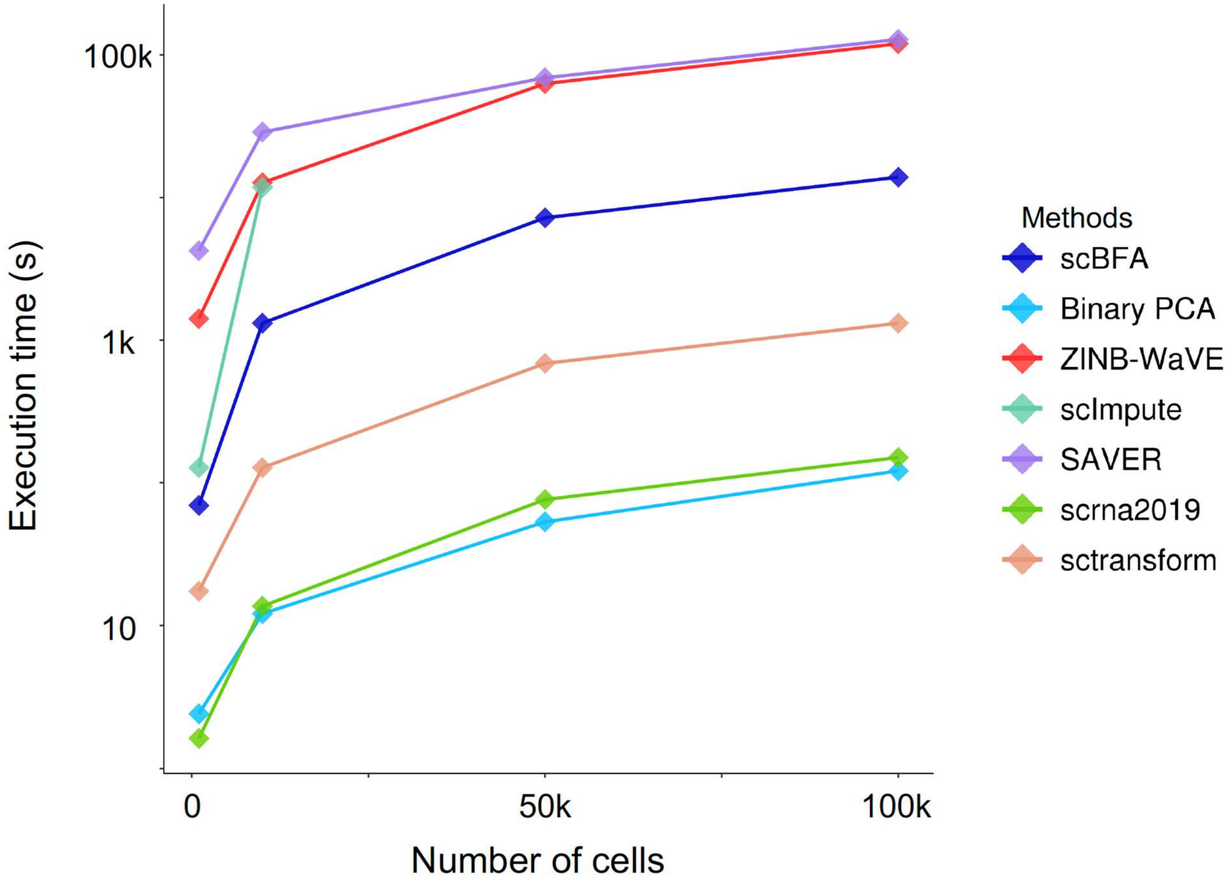

**Supplementary Figure 25: Fast scBFA approximation is one of the fastest dimensionality reduction methods.** Execution time of gene detection models (scBFA and Binary PCA) versus quantification models (ZINB-WaVE, scImpute, scrna2019, sctransform, SAVER) on different dataset sizes, as subsampled from the 1.3 million scRNA-seq mouse brain dataset generated from 10x Genomics.

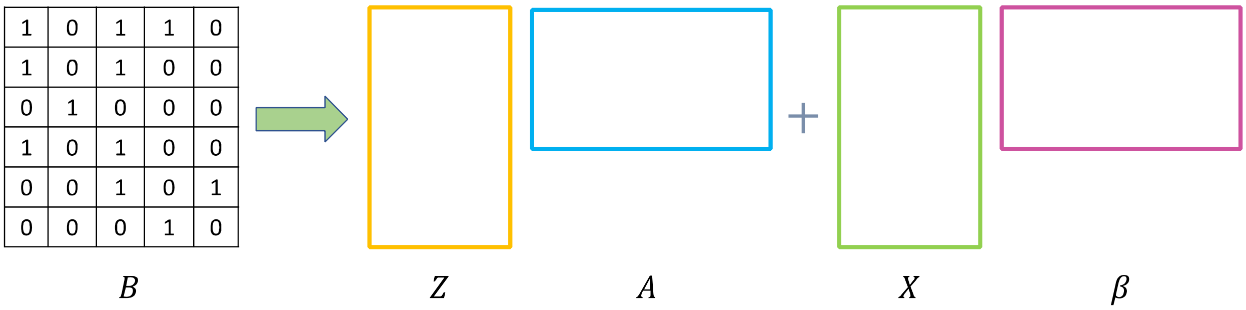

**Supplementary Figure 26: Schematic of scBFA.** $B$ is a matrix of size $N$ (number of cells) by $G$ (number of genes) encoding the gene detection pattern of the input data, where $B_{ij} = 0$ if no molecule (UMI or read) maps to gene $i$ in cell $j$, otherwise $B_{ij} = 1$. BFA decomposes $B$ into a $K$-dimensional embedding matrix $Z$ (factor scores) and a loading matrix $A$. An optional observed cell covariate matrix $X$ is used to model batch effects and other nuisance cell-wise factors. $\beta$ is the corresponding coefficient matrix of $X$.

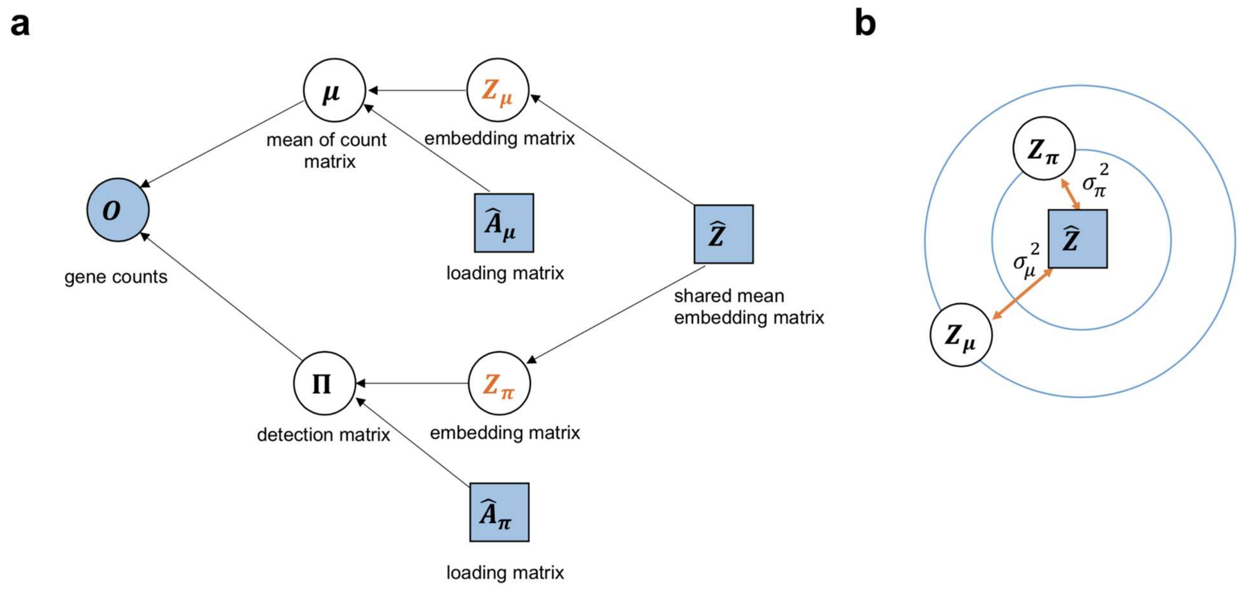

**Supplementary Figure 27: The main components of the generative process of the scRNA-seq simulation framework.** (**a**) Given $N$ cells $(i = 1, \dots, N)$ and $G$ genes $(j = 1, \dots, G)$, our simulation ultimately seeks to generate an observed gene count matrix $\boldsymbol{O}$. $\boldsymbol{O}$ depends on two internal latent variables: a matrix $\boldsymbol{\mu}$ representing gene counts, and a matrix $\boldsymbol{\Pi}$ representing the gene detection pattern. If $\boldsymbol{\Pi}_{ij} = 1$, then $\boldsymbol{O}_{ij} = 0$. Otherwise, if $\boldsymbol{\Pi}_{ij} = 0$, then $\boldsymbol{O}_{ij}$ is sampled from a negative binomial distribution whose mean is $\mu_{ij}$. $\boldsymbol{\mu}$ is determined by the product of a count-based embedding space $\boldsymbol{Z}_{\boldsymbol{\mu}}$ and loading matrix $\widehat{\boldsymbol{A}}_{\boldsymbol{\mu}}$, while $\boldsymbol{\Pi}$ is sampled from a Bernoulli distribution that is parameterized through the product of a separate detection-based embedding $\boldsymbol{Z}_{\boldsymbol{\pi}}$ and loading $\widehat{\boldsymbol{A}}_{\boldsymbol{\pi}}$. $\widehat{\boldsymbol{A}}_{\boldsymbol{\mu}}$ and $\widehat{\boldsymbol{A}}_{\boldsymbol{\pi}}$ are parameters that are estimated prior to the simulation by applying ZINB-WaVE to a real dataset. The count and detection-based embeddings $\boldsymbol{Z}_{\boldsymbol{\mu}}$ and $\boldsymbol{Z}_{\boldsymbol{\pi}}$ respectively are sampled from a distribution with the same mean $\widehat{\boldsymbol{Z}}$ (also learned from applying ZINB-WaVE to real data) but with separate variance terms $\sigma_\mu^2$ and $\sigma_\pi^2$, respectively. These separate variance terms allow us to simulate different noise levels for detection and quantification. (**b**) Graphical illustration indicating that $\boldsymbol{Z}_{\boldsymbol{\mu}}$ and $\boldsymbol{Z}_{\boldsymbol{\pi}}$ are sampled from a common distribution centered on $\widehat{\boldsymbol{Z}}$, though with separate variance terms $\sigma_\mu^2$ and $\sigma_\pi^2$, respectively.

**Supplementary Table 1: Summary of scRNA-seq cell type identification benchmark datasets.**

| Dataset | # of cell types | # of cells | Year | Protocol | Source |
|---|---|---|---|---|---|
| Dendritic | 15 | 1378 | 2014 | full length | http://imlspenticton.uzh.ch/robinson_lab/conquer/data-mae/GSE48968-GPL13112.rds |
| MGE | 5 | 1124 | 2017 | full length | https://www.ncbi.nlm.nih.gov/geo/query/acc.cgi?acc=GSE104157 |
| HSCs | 6 | 949 | 2017 | full length | https://www.ncbi.nlm.nih.gov/geo/query/acc.cgi?acc=GSE100426 |
| Intestinal | 4 | 2891 | 2015 | UMI | http://imlspenticton.uzh.ch/robinson_lab/conquer/data-mae/GSE62270-GPL17021.rds |
| MEM-T | 7 | 2244 | 2018 | UMI | https://www.ncbi.nlm.nih.gov/geo/query/acc.cgi?acc=GSE106540 |
| Pancreatic | 5 | 1152 | 2016 | UMI | http://imlspenticton.uzh.ch/robinson_lab/conquer/data-mae/GSE81076-GPL18573.rds |
| PBMC | 10 | 7666 | 2017 | UMI | https://www.ncbi.nlm.nih.gov/geo/query/acc.cgi?acc=GSE100866 |
| DC | 4 | 950 | 2016 | full length | https://www.ncbi.nlm.nih.gov/geo/query/acc.cgi?acc=GSE89232 |
| Myeloid | 5 | 1816 | 2018 | full length | https://www.ncbi.nlm.nih.gov/geo/query/acc.cgi?acc=GSE123025 |
| H7-ESC | 9 | 651 | 2016 | full length | http://imlspenticton.uzh.ch/robinson_lab/conquer/data-mae/SRP073808.rds |
| mESCs | 3 | 288 | 2015 | full length | http://imlspenticton.uzh.ch/robinson_lab/conquer/data-mae/EMTAB2805.rds |
| LPS | 5 | 839 | 2017 | full length | http://imlspenticton.uzh.ch/robinson_lab/conquer/data-mae/GSE94383.rds |
| LSK | 4 | 1188 | 2017 | UMI | https://www.ncbi.nlm.nih.gov/geo/query/acc.cgi?acc=GSE100037 |
| HSPC | 3 | 1826 | 2016 | full length | https://www.ncbi.nlm.nih.gov/geo/query/acc.cgi?acc=GSE81682 |

**Supplementary Table 2: Summary of gene intersection between the top 2,000 genes selected by HVG and HEG**

| Dataset | # genes in intersection between HVG and HEG |
|---|---|
| Dendritic | 345 |
| MGE | 293 |
| HSCs | 906 |
| Intestinal | 490 |
| MEM-T | 383 |
| Pancreatic | 691 |
| PBMC | 582 |
| DC | 301 |
| Myeloid | 398 |
| H7-ESC | 185 |
| mESCs | 227 |
| LPS | 260 |
| LSK | 613 |
| HSPC | 60 |

**Supplementary Table 3: Cell surface marker list for the PBMC benchmark.**

| Marker Genes | Source |
|---|---|
| CD3D, CD8A, NKG7, FCER1A, CD16, S100A8, S100A9, CD79A, CD79b, CD4, CCR10, PF4, TNFRSF18 | https://www.nature.com/articles/ncomms14049 |
| CD94, GATA3 (Important regulator), NKG2 | https://link.springer.com/content/pdf/10.1385%2FIR%3A35%3A3%3A263.pdf |
| CD163, GZMB, FCRL2, CD40LG, CRTAM, SIGLEC10, FOXP3, IL17A, CXCR5, KLRC1, PRF1, LINC00926, RP11-291B21.2 | https://www.ncbi.nlm.nih.gov/pmc/articles/PMC1895851/ |
| GZMB, HLA_DPB1, DUSP5, LY96, NOD2, STAM, CTSW, CD160, CD244, NCR3 | http://journals.plos.org/plosbiology/article?id=10.1371/journal.pbio.1001148 |
| CD3G, LEF1, TCF7 | https://www.ncbi.nlm.nih.gov/pubmed/16424171 |
| CD8B | https://www.ncbi.nlm.nih.gov/pmc/articles/PMC4686144/ |

**Supplementary Table 4: Cell surface marker list for the HSC benchmark.**

| Marker Genes | Source |
|---|---|
| Tnf, Irf1, Tlr2, Cxcl2, Traf1, Ccl5, Cxcl1, Clec4e, Saa3, Il1a, Il1b, Mmp13, Il10, Lcn2, Il12b, Nos2, Ifitm3 | https://www.ncbi.nlm.nih.gov/pubmed/30540934 |
| Gfi1b, Selp, Klf1, Pf4, Gp9, Zfpm1, Ikzf1, Ikzf2, Flt3, Il7ra | http://www.bloodjournal.org/content/121/22/4463?sso-checked=true |
| Tie2 | https://www.ncbi.nlm.nih.gov/pubmed/15260986 |
| Alcam | https://www.ncbi.nlm.nih.gov/pubmed/23280653 |
| Kit | http://jem.rupress.org/content/211/2/217 |
| Slamf1 | https://www.ncbi.nlm.nih.gov/pmc/articles/PMC2851806/ |
| Matk/Chk | https://www.ncbi.nlm.nih.gov/pmc/articles/PMC1895851/ |
| Cd81 | http://journals.plos.org/plosbiology/article?id=10.1371/journal.pbio.1001148 |
| Ccl5 | http://www.bloodjournal.org/content/bloodjournal/119/11/2500.full.pdf?sso-checked=true |
| Runx1, Notch1, Meis1 | https://link.springer.com/chapter/10.1007%2F978-94-007-6621-1_11 |
| Ly6a | https://www.ncbi.nlm.nih.gov/pubmed/22136929 |
| Egr1 | https://www.sciencedirect.com/science/article/pii/S1934590908000568 |
| Mecom | https://www.ncbi.nlm.nih.gov/pubmed/21666053 |
| Vwf | https://www.ncbi.nlm.nih.gov/pmc/articles/PMC3879699/ |
| Lmo2 | https://www.ncbi.nlm.nih.gov/pmc/articles/PMC3092146 |
| Igta2b | https://journals.plos.org/plosone/article?id=10.1371/journal.pone.0043300 |
| Flt3 | http://journals.plos.org/plosone/article?id=10.1371/journal.pone.0138257 |
| Gata2 | https://www.ncbi.nlm.nih.gov/pmc/articles/PMC4797020/ |
| Gfi1, Gfi1b | https://www.ncbi.nlm.nih.gov/pmc/articles/PMC524350/ |
| Meis1 | https://www.ncbi.nlm.nih.gov/pmc/articles/PMC4795694/ |
| Gata3 | https://www.pnas.org/content/102/7/2448 |
| Ly6a | https://www.ncbi.nlm.nih.gov/pubmed/22136929 |

**Supplementary Table 5: Cell surface marker list for the Pancreatic benchmark.**

| Marker Genes | Source |
|---|---|
| GCG, KRT19, PRSS1, SST, GCG, INS, PPY, FTH1, KRT7, PRSS2 | https://www.cell.com/cell-stem-cell/fulltext/S1934-5909(16)30094-7?_returnURL=https%3A%2F%2Flinkinghub.elsevier.com%2Fretrieve%2Fpii%2FS1934590916300947%3Fshowall%3Dtrue |
| IRX2, GC, LOXL4, SPP1, IAPP, PNLIP, PRG4, IAPP, ALDH1A1, PCSK2 | https://www.cell.com/cell-systems/pdf/S2405-4712(16)30292-7.pdf |
| TSPAN7, FXYD2, TMEM27, SEZ6L2, LRP11, DISP2, DDR1, DNER, NPTX2 | https://link.springer.com/article/10.1007%2Fs00125-011-2295-1 |
| SCG5, GAD2, CPB1, CELA3B, SYCN, GATM, SLC4A4, DPEP1 | https://www.ncbi.nlm.nih.gov/pmc/articles/PMC4278897/ |
| SCGN | https://www.ncbi.nlm.nih.gov/pmc/articles/PMC5510001/ |
| AMY2A, CDKN1C | https://www.ncbi.nlm.nih.gov/pmc/articles/PMC3582140/ |
| CFTR, GP2, CPA1, CEL, PNLIP | https://www.ncbi.nlm.nih.gov/pmc/articles/PMC5494890/ |
| TFF1 | https://www.ncbi.nlm.nih.gov/pmc/articles/PMC4319540/ |
| SPRINK1 | https://www.ncbi.nlm.nih.gov/pmc/articles/PMC3097947/ |
| AQP8 | https://www.ncbi.nlm.nih.gov/pubmed/11254497 |
| CLDN10 | https://www.ncbi.nlm.nih.gov/pmc/articles/PMC3288608/ |
| CHGA | https://viacyte.com/wp-content/uploads/Kelly_Nature_Biotech_07_2011.pdf |
| PECAM1 | http://jcs.biologists.org/content/118/18/4103 |
| MMP2 | https://www.ncbi.nlm.nih.gov/pubmed/15734845 |
| C3 | https://www.cell.com/cell-metabolism/pdfExtended/S1550-4131(18)30574-6 |
| REG1A | https://bmcendocrdisord.biomedcentral.com/articles/10.1186/1472-6823-12-13 |

**Supplementary Table 6: Group I and Group II benchmark list.**

| Dataset | Performance | Gene Selection |
|---|---|---|
| Dendritic | Group I | HEG |
| MGE | Group I | HEG |
| HSCs | Group I | HEG |
| Intestinal | Group I | HEG |
| MEM-T | Group I | HEG |
| Pancreatic | Group I | HEG |
| PBMC | Group I | HEG |
| DC | Group I | HEG |
| Myeloid | Group I | HEG |
| H7-ESC | Group II | HEG |
| mESCs | Group II | HEG |
| LPS | Group II | HEG |
| LSK | Group II | HEG |
| HSPC | Group II | HEG |
| Dendritic | Group I | HVG |
| MGE | Group I | HVG |
| HSCs | Group I | HVG |
| Intestinal | Group I | HVG |
| MEM-T | Group I | HVG |
| Pancreatic | Group I | HVG |
| PBMC | Group I | HVG |
| DC | Group I | HVG |
| Myeloid | Group I | HVG |
| H7-ESC | Group I | HVG |
| mESCs | Group I | HVG |
| LPS | Group I | HVG |
| LSK | Group II | HVG |
| HSPC | Group II | HVG |

**Supplementary Table 7: Summary of representative scRNA-seq datasets.**

| ID | Year | Protocol | Source |
|---|---|---|---|
| GSE101601 | 2017 | UMI | https://www.ncbi.nlm.nih.gov/geo/query/acc.cgi?acc=GSE101601 |
| GSE106707 | 2017 | UMI | https://www.ncbi.nlm.nih.gov/geo/query/acc.cgi?acc=GSE106707 |
| GSE110558 | 2018 | UMI | https://www.ncbi.nlm.nih.gov/geo/query/acc.cgi?acc=GSE110558 |
| GSE110692 | 2018 | UMI | https://www.ncbi.nlm.nih.gov/geo/query/acc.cgi?acc=GSE110692 |
| GSE119097 | 2018 | UMI | https://www.ncbi.nlm.nih.gov/geo/query/acc.cgi?acc=GSE119097 |
| GSE56638 | 2014 | full length | https://www.ncbi.nlm.nih.gov/geo/query/acc.cgi?acc=GSE56638 |
| GSE72056 | 2015 | full length | https://www.ncbi.nlm.nih.gov/geo/query/acc.cgi?acc=GSE72056 |
| GSE81682 | 2016 | full length | https://www.ncbi.nlm.nih.gov/geo/query/acc.cgi?acc=GSE81682 |
| GSE85527 | 2016 | UMI | https://www.ncbi.nlm.nih.gov/geo/query/acc.cgi?acc=GSE85527 |
| GSE86977 | 2016 | full length | https://www.ncbi.nlm.nih.gov/geo/query/acc.cgi?acc=GSE86977 |
| GSE95432 | 2017 | full length | https://www.ncbi.nlm.nih.gov/geo/query/acc.cgi?acc=GSE95432 |
| GSE98816 | 2017 | full length | https://www.ncbi.nlm.nih.gov/geo/query/acc.cgi?acc=GSE98816 |
| GSE95315 | 2017 | UMI | https://www.ncbi.nlm.nih.gov/geo/query/acc.cgi?acc=GSE95315 |
| GSE95752 | 2017 | full length | https://www.ncbi.nlm.nih.gov/geo/query/acc.cgi?acc=GSE95752 |
| GSE76381 | 2016 | UMI | https://www.ncbi.nlm.nih.gov/geo/query/acc.cgi?acc=GSE76381 |
| GSE110679 | 2018 | UMI | https://www.ncbi.nlm.nih.gov/geo/query/acc.cgi?acc=GSE110679 |
| GSE99888 | 2018 | UMI | https://www.ncbi.nlm.nih.gov/geo/query/acc.cgi?acc=GSE99888 |
| GSE52529 | 2014 | full length | http://imlspenticton.uzh.ch/robinson_lab/conquer/data-mae/GSE52529-GPL16791.rds |
| GSE60749 | 2014 | full length | http://imlspenticton.uzh.ch/robinson_lab/conquer/data-mae/GSE60749-GPL13112.rds |
| GSE63818 | 2015 | full length | http://imlspenticton.uzh.ch/robinson_lab/conquer/data-mae/GSE63818-GPL16791.rds |
| GSE71982 | 2015 | full length | http://imlspenticton.uzh.ch/robinson_lab/conquer/data-mae/GSE71982.rds |
| GSE57872 | 2014 | full length | http://imlspenticton.uzh.ch/robinson_lab/conquer/data-mae/GSE57872.rds |
| GSE102299 | 2017 | UMI | https://www.ncbi.nlm.nih.gov/geo/query/acc.cgi?acc=GSE102299 |
| GSE48968 | 2014 | full length | http://imlspenticton.uzh.ch/robinson_lab/conquer/data-mae/GSE48968-GPL13112.rds |
| GSE104157 | 2017 | full length | https://www.ncbi.nlm.nih.gov/geo/query/acc.cgi?acc=GSE104157 |
| GSE100426 | 2017 | full length | https://www.ncbi.nlm.nih.gov/geo/query/acc.cgi?acc=GSE100426 |
| GSE62270 | 2015 | UMI | http://imlspenticton.uzh.ch/robinson_lab/conquer/data-mae/GSE62270-GPL17021.rds |
| GSE106540 | 2018 | UMI | https://www.ncbi.nlm.nih.gov/geo/query/acc.cgi?acc=GSE106540 |

| | | | |
|---|---|---|---|
| GSE81076 | 2016 | UMI | http://imlspenticton.uzh.ch/robinson_lab/conquer/data-mae/GSE81076-GPL18573.rds |
| SRP073808 | 2016 | full length | http://imlspenticton.uzh.ch/robinson_lab/conquer/data-mae/SRP073808.rds |
| EMTAB2805 | 2015 | full length | http://imlspenticton.uzh.ch/robinson_lab/conquer/data-mae/EMTAB2805.rds |
| GSE94383 | 2017 | full length | http://imlspenticton.uzh.ch/robinson_lab/conquer/data-mae/GSE94383.rds |

**Supplementary Table 8: Summary of benchmarks used in trajectory inference obtained from** https://zenodo.org/record/1443566#.XP86LtNKiRc

| Dataset | Alias |
|---|---|
| aging-hsc-old_kowalczyk.rds | HSC-old |
| aging-hsc-young_kowalczyk.rds | HSC-young |
| cell-cycle_buettner.rds | mESC |
| developing-dendritic-cells_schlitzer.rds | Develop-dc |
| germline-human-female-weeks_li.rds | F-embryo |
| germline-human-male-weeks_li.rds | M-embryo |
| hematopoiesis-gates_olsson.rds | HSC-gate |
| human-embryos_petropoulos.rds | H-embryo |
| myoblast-differentiation_trapnell.rds | Myoblast |
| NKT-differentiation_engel.rds | Pan-alpha |
| pancreatic-alpha-cell-maturation_zhang.rds | NKT |
| psc-astrocyte-maturation-glia_sloan.rds | Glia |
| psc-astrocyte-maturation-neuron_sloan.rds | Neuron |
| stimulated-dendritic-cells-LPS_shalek.rds | LPS-DC |
| stimulated-dendritic-cells-PAM_shalek.rds | PAM-DC |
| stimulated-dendritic-cells-PIC_shalek.rds | PIC-DC |
| pancreatic-beta-cell-maturation_zhang.rds | Pan-beta |
| mesoderm-development_loh.rds | mesoderm |
